# Wake EEG oscillation dynamics reflect both sleep pressure and brain maturation across childhood and adolescence

**DOI:** 10.1101/2024.02.24.581878

**Authors:** Sophia Snipes, Valeria Jaramillo, Elena Krugliakova, Carina Volk, Melanie Furrer, Mirjam Studler, Monique LeBourgeois, Salome Kurth, Oskar G. Jenni, Reto Huber

**Affiliations:** Child Development Center, University Children’s Hospital Zurich, University of Zurich, Zurich, Switzerland; Paris Brain Institute, Sorbonne Université, Inserm-CNRS, Paris, France; School of Psychology, University of Surrey, Guildford, UK; Surrey Sleep Research Centre, Faculty of Health and Medical Sciences, University of Surrey, Guildford, United Kingdom; UK Dementia Research Institute Care Research and Technology Centre, Imperial College London and the University of Surrey, Guildford, UK; Donders Institute, Radboud University Medical Center, Nijmegen, the Netherlands; Center for MR Research, University Children’s Hospital Zurich, Switzerland; Department of Social Neuroscience and Social Psychology, Institute of Psychology, University of Bern, Bern, Switzerland; University of Colorado at Boulder, Department of Integrative Physiology, Boulder, Colorado, USA; The Warren Alpert Medical School of Brown University, Department of Psychiatry and Human Behavior, Providence, Rhode Island, USA; In memoriam; Department of Psychology, University of Fribourg, Fribourg, Switzerland; Department of Pulmonology, University Hospital Zurich, Zurich, Switzerland; Department of Child and Adolescent Psychiatry and Psychotherapy, Psychiatric Hospital, University of Zurich, Switzerland

**Author notes:** Corresponding author: Sophia Snipes.

**Keywords:** development, EEG, sleep, synaptic homeostasis hypothesis, aperiodic activity, oscillation bursts

## Abstract

An objective measure of brain maturation is highly insightful for monitoring both typical and atypical development. Slow wave activity, recorded in the sleep electroencephalogram (EEG), reliably indexes age-related changes in sleep pressure as well as deficits related to developmental disorders such as attention-deficit hyperactivity disorder (ADHD). We aimed to determine whether wake EEG measured before and after sleep could index the same developmental changes in sleep pressure, using data collected from 163 participants 3-25 years old. We analyzed age- and sleep-dependent changes in two measures of oscillatory activity, amplitudes and density, as well as two measures of aperiodic activity, offsets and exponents. We then compared these wake measures to sleep slow wave amplitudes and slopes. Finally, we compared wake EEG in children with ADHD (N=58) to neurotypical controls. Of the four wake measures, only oscillation amplitudes consistently exhibited the same changes as sleep slow waves. Wake amplitudes decreased with age, decreased after sleep, and this overnight decrease decreased with age. Furthermore, wake amplitudes were significantly related to both sleep slow wave amplitudes and slopes. Wake oscillation densities decreased overnight in children but increased overnight in adolescents and adults. Aperiodic offsets decreased linearly with age, decreased after sleep, and were significantly related to sleep slow wave amplitudes. Aperiodic exponents also decreased with age, but increased after sleep. No wake measure showed significant effects of ADHD. Overall, our results indicate that wake oscillation amplitudes, and to some extent aperiodic offsets, behave like sleep slow waves across sleep and development. At the same time, overnight changes in oscillation densities independently reflect some yet-unknown shift in neural activity around puberty.

## 1 INTRODUCTION

The EEG is one of few tools available to study the human developing brain already from birth (Korotchikova et al., 2009). It is non-invasive, relatively cheap, and provides a realtime readout of neuronal activity. It is an incredibly rich signal, with the potential as a prognostic and diagnostic tool for both typical development and disease. Sleep EEG, and in particular slow wave activity (0.5-4 Hz) during NREM sleep, has proven especially sensitive to brain maturation (Campbell & Feinberg, 2009) and developmental disorders such as ADHD (Furrer et al., 2019). This is because slow wave activity reflects the overall synchronicity of the brain, which decreases with age following decreasing synaptic density across adolescence (Campbell & Feinberg, 2009; Huttenlocher, 1979; Jenni & Carskadon, 2004), and may be lower in ADHD due to reduced cortical thickness (Shaw et al., 2006). Furthermore, slow wave activity is greater in occipital regions in younger children, and greater in frontal regions in adolescents and adults (Kurth et al., 2010), possibly reflecting the slower maturation of higher order association areas (Shaw et al., 2008).

In addition to these developmental phenomena, sleep slow waves reflect the buildup and dissipation of homeostatic sleep pressure, increasing following wake and decreasing during sleep (Borbély, 1982). These hourly changes in sleep pressure detected through slow wave activity are hypothesized to reflect synaptic *plasticity*: synaptic strength increases with wake and daytime learning, and decreases during sleep (Tononi & Cirelli, 2003, 2014). Generally, plasticity decreases with brain maturation across childhood and adolescence and this is reflected in decreases in the overnight *changes* in slow waves with age (Jaramillo et al., 2020). Thus, both absolute slow wave activity and changes in slow wave activity are markers of brain development, reflecting average synaptic density and synaptic plasticity respectively. While these predictions have been repeatedly tested and validated in sleep, their consequences on the wake EEG have not been as systematically investigated. Previous research has found correlations between wake and sleep EEG power (Finelli et al., 2000), however, greater insights can be derived from more specific analyses.

The EEG is made up of both *periodic* activity and *aperiodic* activity (Figure 1A, Donoghue et al., 2020). Periodic activity refers to oscillations, which appear as quasi-gaussian bumps in the power spectrum at their corresponding frequency. Instead, aperiodic activity is a form of background “noise,” producing the characteristic 1/f curve in the EEG power spectrum. Aperiodic activity is defined by its *offset* (reflecting the overall aperiodic power) and *exponent* (the steepness of the curve). Changes in exponent in particular are hypothesized to reflect alterations in excitatory/inhibitory balance of neuronal activity (Brake et al., 2024; Gao et al., 2017), and this could explain the multitude of conditions which affect aperiodic signals.

**Figure 1:**
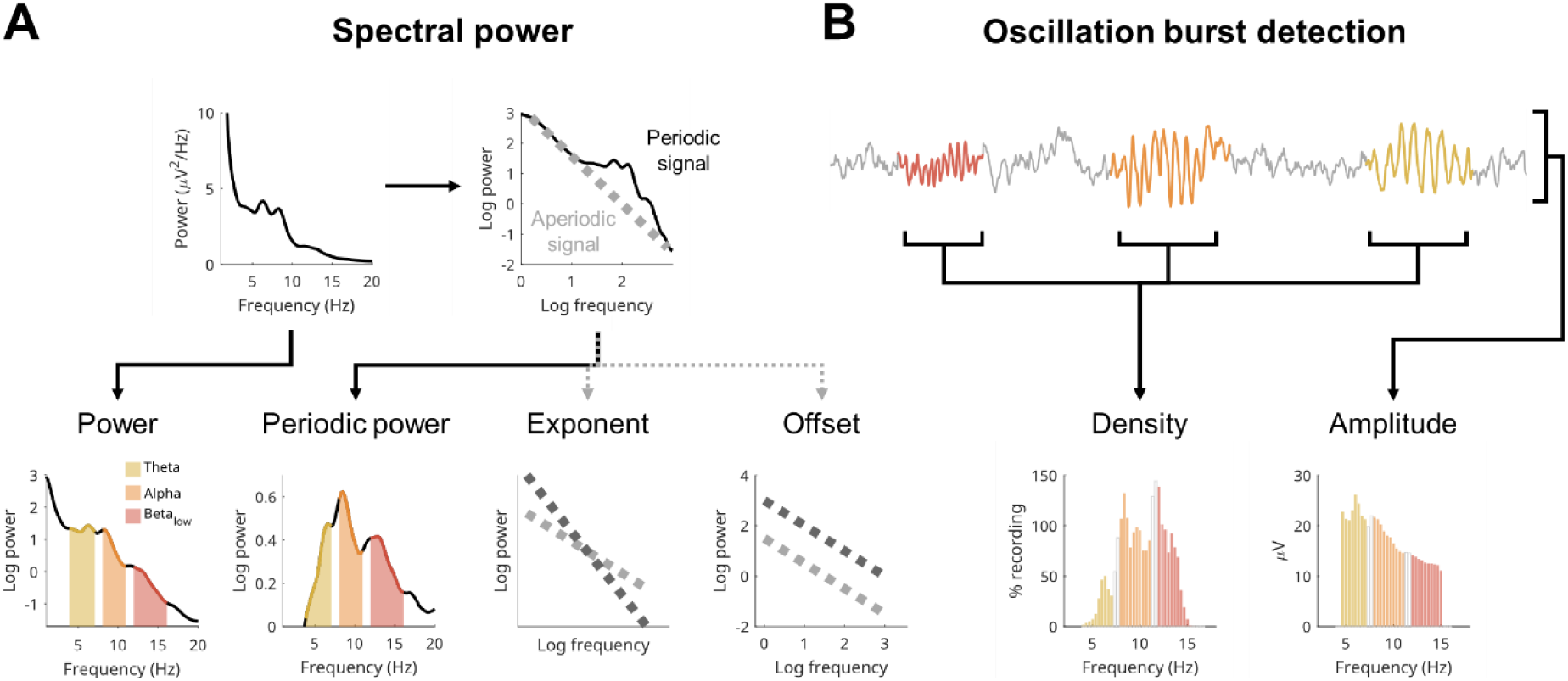
Wake EEG measures. **A**: Measures based on spectral power. Given the nature of power, it is traditionally analyzed log-transformed to have more normally distributed values. These are then aggregated into bands. Here, we focus on classical wake EEG bands: theta (4-7 Hz, yellow), alpha (8-11 Hz, orange) and low beta (12-16 Hz, red), with gaps between bands to avoid overlapping information. The example comes from a 15-year-old male participant, used for the entire figure. The EEG signal is composed of aperiodic “background activity” (gray signal in B) and oscillatory activity (colored signal in B). When plotting the power spectrum on a log-log scale, a line can be fitted to the aperiodic component of the signal, which can be subtracted from the whole spectrum, leaving behind only periodic power. The aperiodic line can then be quantified by its offset (where it intersects 0 on the log-log scale, i.e. the log power at 1 Hz), and its exponent (how tilted it is). Thus, the power spectrum provides four measures: log-transformed power, periodic power, exponent, and offset of the aperiodic signal. **B**: Cycle-by-cycle analysis is used to detect bursts of oscillations by identifying sections of the EEG signal that show periodic activity (colored), relative to the aperiodic background activity (gray). Once bursts are detected, there are two main parameters to quantify them: density (how much of the signal in time contains an oscillation) and amplitude (average peak-to-peak voltage of each oscillation). Densities are expressed in percentage, and when pooling bursts detected in all channels, they can easily exceed 100%, as the same burst will appear in multiple channels and multiple bursts can co-occur. Amplitudes are in microvolts. The EEG trace was stitched together for illustrative purposes. N.B. Beta periodic power is lower than alpha periodic power (in panel A), but their densities (in B) are roughly the same; this is because periodic power is also influenced by the lower amplitudes of low beta compared to alpha. The densities of other participants are provided in Suppl. Figure 1-1.

Aperiodic exponents reflect levels of consciousness, becoming progressively steeper with increasing sleep depth (Lendner et al., 2020, 2023; Schneider et al., 2022), anesthesia (Colombo et al., 2019), and disorders of consciousness (Colombo et al., 2023; Maschke et al., 2023). Additionally, exponents become progressively shallower during sleep, reflecting the dissipation of sleep pressure (Bódizs et al., 2024; Horváth et al., 2022). Brain maturation also strongly affects exponents and offsets, becoming shallower and lower from childhood to adulthood (Cellier et al., 2021; Hill et al., 2022; Tröndle et al., 2022), with age and sleep depth interacting (Favaro et al., 2023). Lastly, differences in aperiodic exponents and offsets have been found in children with ADHD compared to neurotypical controls (Ostlund et al., 2021; Robertson et al., 2019).

Periodic activity, like aperiodic activity, can be estimated from spectral power by simply subtracting the aperiodic activity from overall power, giving *periodic power* (Figure 1A). However this misses two important independent changes that can happen to oscillations: they can change in *amplitude*, and they can change in *density* (Figure 1B). The amplitudes of oscillations reflect the synchronicity of the oscillating neuronal population, and that synchronicity is determined both by the number of neurons in phase with each other and the strength of their synaptic connections. Whether an oscillation occurs at all (i.e., density) will instead depend on the activity of “pacemaker” interneurons which entrain a population of neurons to the same rhythm (Le Bon-Jego & Yuste, 2007; Perkel et al., 1964), and this pattern of rhythmic firing will be in service of some underlying function that will come and go as needed. In short, amplitudes reflect synchronicity and densities reflect activity.

Since oscillation amplitudes reflect synchronicity, this means they should reflect the same information as slow waves measured during sleep. Sleep slow wave activity increases along a saturating exponential function relative to the prior duration of wakefulness, increasing rapidly after only a few hours awake and increasing less and less with further hours awake (Borbély, 1982; Dijk et al., 1987). We found that wake oscillation amplitudes follow the same trajectory across 24 hours awake (Snipes et al., 2023). Instead, alpha oscillation densities (8-12 Hz) decreased, supporting the specificity of the effect to amplitudes and masking the effect when measuring spectral power.

Given these results, we hypothesized that wake oscillation amplitudes should behave like sleep slow waves also across development: absolute amplitudes should decrease with age, and overnight changes in amplitude should decrease with age. Likewise, changes in amplitude should also manifest an anterior-posterior gradient with age: larger amplitudes in occipital regions in children and larger in frontal regions in adults. Furthermore, given that children with ADHD have lower sleep slow wave activity, they should likewise have lower wake amplitudes. In short, if both wake amplitudes and sleep slow waves are supposed to reflect brain-wide synchronicity, then wake amplitudes should also reflect sleep pressure, brain development, and the pathophysiology of ADHD. We further hypothesized that all these effects would be specific to amplitudes, with other wake EEG parameters changing independently with sleep and age.

Finally, we expected that wake amplitudes, more than any other wake EEG measure, should directly correlate with specific sleep markers of synaptic plasticity: slow wave amplitudes and slopes (the steepness of the waves) measured at the beginning and end of the night (Esser et al., 2007; Jaramillo et al., 2020; Riedner et al., 2007; Vyazovskiy et al., 2009). Slopes in particular are thought to reflect changes in synaptic strength. The steepness of a slow wave is determined by the rate at which neuronal populations synchronize and desynchronize firing, and increased synaptic strength will increase the speed of such synchronization (Vyazovskiy et al., 2009). Therefore, any wake measure of sleep pressure and synaptic plasticity should correlate with both sleep slow wave amplitudes and especially sleep slow wave slopes.

To answer these questions, we analyzed data collected from previous studies at the University Children’s Hospital of Zurich, with high-density wake EEG recordings measured the evening before and morning after a night of sleep. The final dataset included 105 neurotypical participants from the ages of 3.5 to 25, and 58 children with ADHD. From 44 neurotypical participants, the wake EEG was further compared to their sleep EEG.

## 2 METHODS

### 2.1 Datasets

The data for this manuscript was assembled from previous studies conducted between 2008 and 2021. The participant demographics of each dataset are in Table 1. In total, we included 163 participants between the ages 3.5 and 24.7, 38% female, 7% lefthanded. Of these, 36% were diagnosed with ADHD at the department for Child and Adolescent Psychiatry at the University of Zurich, the outpatient clinic of the Child Development Center, and at private children’s clinics in Zurich Oerlikon. Patients were not excluded based on medication status, and therefore were a mixture of medicated, previously medicated, and unmedicated (see Table 2, and (Furrer et al., 2019; Ringli et al., 2013)). Otherwise, all participants were screened by telephone such that they all were completely healthy, took no (other) medication, had no (other) comorbidities, and were good sleepers. All participants were recruited from canton Zurich, Switzerland, and recorded at the University Children’s Hospital of Zurich, except for the children from 3.5 to 8 (Dataset2009, N=11), who were recruited in Providence, RI, USA, and recorded at home. Sleep time was determined by their individual preferred sleep and wakeup time, which they had to maintain the week prior to each measurement. Wake measurements were done just before going to sleep, and ∼30 minutes after waking up. 115 participants had 2 sessions, spaced at least 1 week apart, both included in these analyses. Depending on the dataset, different paradigms were used involving different wake tasks (described below) and there could be 1-4 tasks at each time point (see Table 1). Every set of tasks for each dataset was repeated in the evening and in the morning. In total, 1243 recordings were included in these analyses lasting on average 4.2 min (task time + buffer − artefacts), with a standard deviation of 2.5 min. The data (averaged across channels) for all participants is provided in Suppl. Data 1.

**Table 1:** All participants’ demographics, split by dataset. The year for each dataset indicates the beginning of data collection. *N* indicates the number of participants. After the mean age of each dataset, the standard deviation is provided in parentheses. *Paradigm* indicates which set of wake tasks were recorded. *Sessions* indicate the number of sessions expected for each dataset, although in practice due to dropouts, some participants only completed 1. *Recordings* indicate the average number of recordings per participant compared to the total number of recordings expected by the experimental paradigm. Recordings were missing either because they contained too many artefacts or were omitted entirely during data collection.

|  | N | Female | Left-handed | ADHD | Age range | Mean age | Paradigm | Sessions | Recordings |
| --- | --- | --- | --- | --- | --- | --- | --- | --- | --- |
|  | # | % | % | % | (years) | (years) |  | # | # |
| *Dataset2008* | 38 | 32 | 0 | 0 | 8.7-23.4 | 14.1 (3.8) | Adaptation | 2 | 7.3/8 |
| *Dataset2009* | 11 | 73 | 0 | 0 | 3.5-8.0 | 5.6 (1.5) | Oddball | 1 | 1.8/2 |
| *Dataset2010* | 28 | 21 | 14 | 100 | 9.7-16.3 | 12.7 (1.9) | Adaptation | 1 | 2.8/4 |
| *Dataset2016* | 18 | 44 | 0 | 0 | 18.4-24.7 | 21.6 (2.1) | Oddball | 2 | 3.9/4 |
| *Dataset2017* | 42 | 43 | 17 | 36 | 8.1-17.6 | 12.2 (2.7) | Attention | 2 | 12.7/16 |
| *Dataset2019* | 26 | 38 | 0 | 58 | 8.8-16.8 | 11.4 (2.0) | Attention | 2 | 10.1/12 |
| *All* | 163 | 38 | 7 | 36 | 3.5-24.7 | 13.2 (4.4) |  |  |  |

**Table 2:** ADHD demographics, split by patient status. For each patient group, the *Oddball* column indicates how many came from a dataset performing the oddball task (rather than the attention tasks). *For 3 patients, medication status was missing, therefore the true total is 58.

|  | N | Female | Left-handed | Age range | Mean age | Oddball |
| --- | --- | --- | --- | --- | --- | --- |
|  | # | % | % | (years) | (years) | % |
| *Medicated in the past* | 6 | 17 | 0 | 9.7-14.8 | 12.5 (2.2) | 33 |
| *Unmedicated* | 14 | 21 | 14 | 9.5-15.3 | 11.3 (1.6) | 64 |
| *Medication the day before* | 22 | 23 | 14 | 8.7-16.3 | 12.3 (2.2) | 45 |
| *Medication the day of* | 13 | 15 | 0 | 9.0-16.1 | 12.3 (2.0) | 54 |
| *All patients* | 55* | 20 | 9 | 8.7-16.3 | 12.1 (2.0) | 51 |
| *Controls* | 105 | 47 | 6 | 3.5-24.7 | 13.8 (5.2) | 58 |

Informed consent was obtained from all adult participants, and from the legal guardians of all children below 14, as well as from adolescent participants 14-18. All studies were approved by the local ethics committees and performed according to the declaration of Helsinki.

#### 2.1.1 Oddball & motor adaptation paradigms

95 participants (Dataset2008, Dataset2009, Dataset2010, Dataset2016) performed an auditory oddball task during their wake EEG. The task lasted 4 minutes and was performed in the evening just before going to bed and in the morning ∼30 minutes after waking up. The task involved 300 tones at ∼80 dB, with an interstimulus interval of 0.8 s. A random 10% of stimuli were targets to which the participant had to push a button in response. For the youngest children (Dataset2009), the 4-minute task was split into 2 segments, and for the adults (Dataset2016) the task was extended to 6 minutes.

66 of these participants (Dataset2008, Dataset2010) also performed a half-hour visuomotor adaptation task (Ghilardi et al., 2000) followed by a second oddball. One dataset (Dataset2008) also included a second session with a control visuomotor task (no adaptation), counterbalanced with the motor adaptation task. The motor tasks were not included in this analysis, because they further differed from evening to morning. For more details on the adaptation task see Wilhelm et al. (2014). The youngest (Dataset2009) and oldest (Dataset2016) participants only conducted one oddball and no motor task. Dataset2016 also had two sessions, one night with phase-targeted auditory stimulation during NREM sleep and the other sham. The experimental conditions did not affect the wake EEG (data not shown).

The sleep data from these participants has been published (Buchmann et al., 2011; Furrer et al., 2019, 2020; Jaramillo et al., 2020; Kurth et al., 2010; Ringli et al., 2013; Volk et al., 2018; Wilhelm et al., 2014), as has a subset of the wake EEG data (Fattinger et al., 2017).

#### 2.1.2 Attention paradigm

68 participants (Dataset2017, Dataset2019) performed three tasks with a focus on attention. These were studies investigating the relationship between slow waves, behavior, and MR spectroscopy (Jaramillo et al., 2020; Volk et al., 2019). This included two sessions to compare the effects of phase targeted auditory stimulation on slow waves in sleep (sham and stimulation; data currently unpublished). The wake tasks were part of the Test Battery for Attentional Performance (TAP) (Zimmermann & Fimm, 2012), which included 2 minutes of a visual Go/No-Go task (respond to 1 stimulus, withhold response to another), 5.5 minutes of a visual and acoustic Alertness task, and then 2 1.5-minute Fixation recordings. For one dataset (Dataset2019) the Go/No-Go task was adapted using cartoon images (extended to 12 minutes), but only 1 Fixation recording was measured (lasting 2 minutes).

### 2.2 EEG recordings and preprocessing

All datasets were measured using 128 channel EGI Geodesic Sensor nets and EGI amplifiers (Electrical Geodesics Inc., Eugene, OR, USA). Wake recordings were done with Cz reference, 1000 Hz sampling rate, and impedances kept below 50 kOhm. All analyses were performed in MATLAB 2023b, with the EEGLAB toolbox v2023.1 (Delorme & Makeig, 2004), the FOOOF/specparam toolbox v1.1.0 (Donoghue et al., 2020), and custom scripts.

EEG data was first mean-centered, then lowpass filtered at 40 Hz (EEGLAB’s *pop_eegfiltnew* function) and notch-filtered at either 50 or 60 Hz (Dataset2009) along with subsequent harmonics. The data was downsampled to 250 Hz, then highpass-filtered over 0.5 Hz (Kaiser filter, stopband = 0.25 Hz, stopband attenuation = 60, passband ripple = 0.05).

Artifacts were removed with a fully automated procedure. Movement and other large artifacts were detected in data filtered between 1 and 40 Hz, in 3 s segments. A segment was labeled a “major artifact” if it exceeded 500 μV, or a “minor artifact” if the correlation with neighboring channels was below .3. Major artifacts were always removed, either by removing all data in all channels during those 3 s, or removing the entire channel with such an artifact, depending on which (channel or segment) removed the least amount of clean data. Minor artifacts were removed in a similar way, removing iteratively either the channel with the most artifactual segments, or the segments with the most artifactual channels, until all channels and all segments had at most 30% of the data containing a minor artifact. Flat channels were removed using EEGLAB’s *clean_artifacts* function. The missing Cz channel was added (as a vector of zeros), then the data was average referenced. Physiological artifacts (blinks, eye movements, muscle tone, heartbeat) were removed with independent component analysis (ICA), with components calculated separately as described in the next section. After these were removed, a second pass was conducted using EEGLAB’s *clean_windows* function (MaxBadChannels = .3, PowerTolerances = [-inf, 12]), then bad segments/channels still containing amplitudes over 140 μV were removed, and finally EEGLAB’s *clean_channels_nolocs* was applied (MinCorrelation = .5, IgnoredQuantile = .1, MaxBrokenTime = .5). Recordings for which more than 25 channels were removed, or which had less than 1 minute of data, were excluded from analysis. In a last step, EEG channels were interpolated, for a total of 123 channels, excluding the external electrodes (49 56, 107, 113) and the face electrodes (126, 127).

#### 2.2.1 Automatic detection and removal of physiological artefacts using ICA

For ICA, a separate copy of the EEG data was preprocessed as previously described, however the data was filtered between 2.5 and 100 Hz, and downsampled to 500 Hz. Automatically detected bad channels and bad time windows were removed, an empty Cz channel added, and then the data was re-referenced to the average of all channels. EEGLAB’s *runica* function was run with principal component analysis (PCA) rank reduction. Then, components were automatically classified with EEGLAB’s *iclabel*, as either brain, muscle, eye, heart, line, channel noise, or other. This function provides a probability score for each label from 0 to 1, so the label with the largest score for each component was taken. Components classified as muscle, eye, or heart were removed. Of the remaining noise classifications (line, channel, other), due to poor classification accuracy, an additional step was implemented. Spectral power was calculated for each component (*pwelch*, 4 s Hanning windows, 50% overlap), then smoothed over 5 Hz (*lowess*) to facilitate model fitting. Unlike for the main analyses, the periodic signal was not of interest, therefore stronger smoothing was possible. The *specparam* algorithm (Donoghue et al., 2020; Ostlund et al., 2022) was applied to the power spectrum between 8 and 30 Hz to extract aperiodic exponents. Components for which the spectral exponent was shallower than 0.5 (so almost flat or even tilted positive), were considered noise and therefore excluded, as they reflected either muscle activity or other nonphysiological signals. Using the manually labeled components in an independent adult dataset (Snipes et al., 2022), we confirmed that this procedure was sufficiently comparable to human detection of artifactual components. We further confirmed that the outcome matched human component classification in a small subset of the children’s data. However, considering the trend towards 0.5 exponents observed in Figure 2, for future datasets with older participants we would recommend a lower threshold.

**Figure 2:**
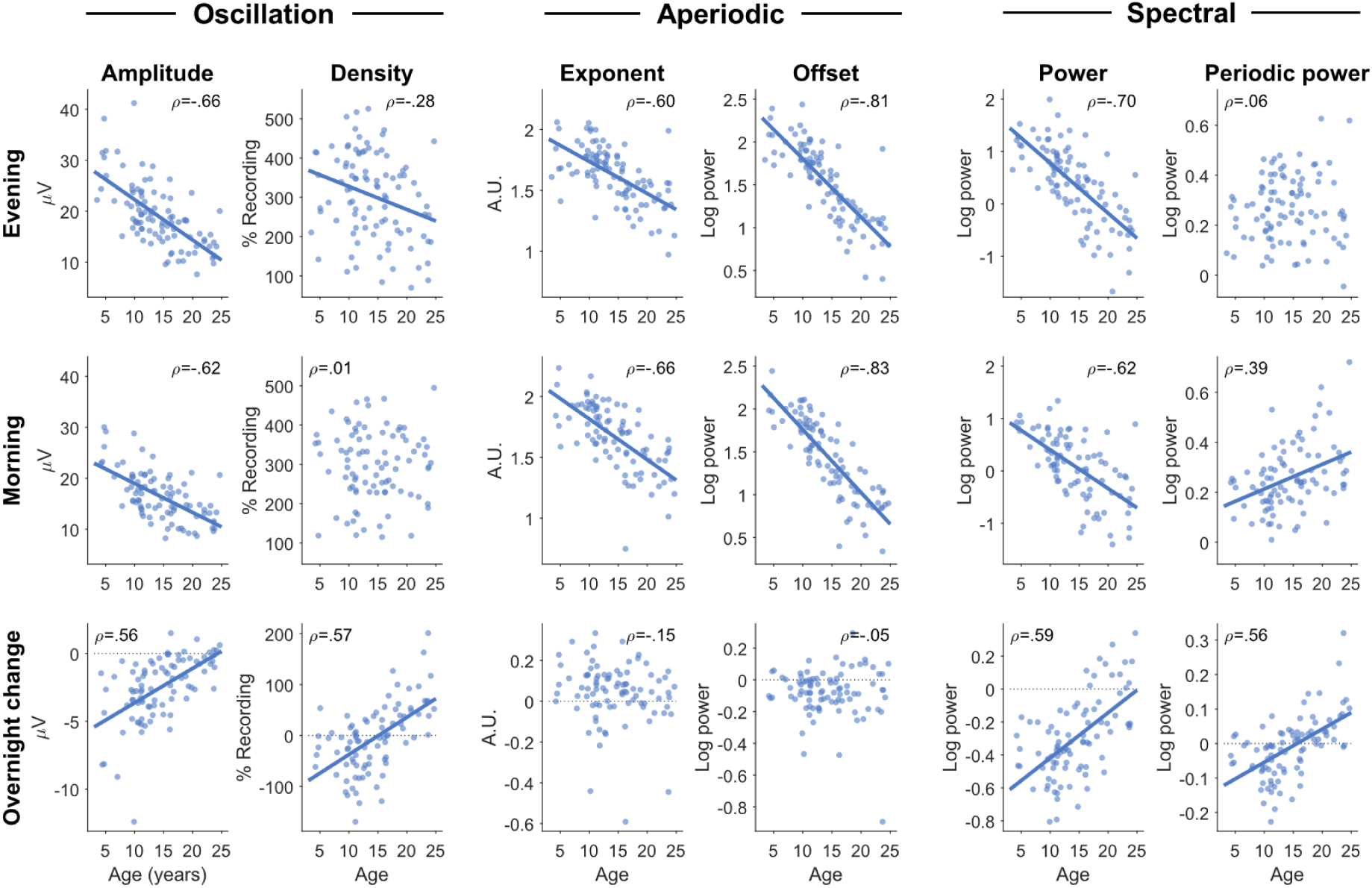
Wake EEG measures correlated with age. Only auditory oddball recordings were included, pooling both neurotypical and ADHD participants. Each dot represents the value for a single participant. For participants with multiple sessions, values across sessions were first averaged. Pearson’s correlations were done for each figure, with rho values provided in the corner. If the p-value was less than .05, a correlation line was drawn (without correcting for multiple comparisons). Amplitudes and densities of oscillations were obtained from burst clusters, pooling all frequencies (4-16 Hz). Exponent values are such that larger values indicate a steeper aperiodic signal; units are “arbitrary” (a.u.), in that exponents are scale-free measures. Power spectra were calculated for each channel, then averaged across all channels, excluding the outermost ring of channels. Power and periodic power were then calculated by averaging values from 4 to 16 Hz. The same figure for the specparam fitting estimates is provided as Suppl. Figure 2-1. The correlations between each measure with the other is provided in Suppl. Figure 2-2.

For the Dataset2009 cohort of <8-year-olds, given how little data there was and how many more movement artifacts, we chose to apply to same manual artifact rejection as in Snipes et al. (2022) to preserve as much data as possible.

### 2.3 Burst detection

Bursts of oscillations were detected using cycle-by-cycle analysis (Cole & Voytek, 2019) implemented in MATLAB (Snipes et al., 2023). Bursts were detected with the same thresholds as in Snipes et al. (2024). Briefly, EEG was narrow-band-pass filtered in overlapping ranges (2-6 Hz, 4-8 Hz…), from which zero-crossings were detected. Then, in the broadband filtered data (0.5-40 Hz), peaks were identified between the zero-crossings, and a cycle was considered an oscillation from positive-to-positive peak. A minimum number of consecutive cycles must meet a set of criteria (monotonicity, period consistency, amplitude consistency, shape consistency, etc.) for this to be considered a burst. Importantly, amplitude itself is never used as a threshold, as this would create a greater dependency between amplitude and density (such that a decrease in an amplitude threshold would result in an automatic increase in density).

Three sets of criteria were used. The first aimed to detect bursts relying on many low-threshold criteria (frequency in range of narrowband filter; PeriodConsistency = .5; AmplitudeConsistency = .4; FlankConsistency = .5; ShapeConsistency = .2; MonotonictyInTime = .4; MonotonicityInAmplitude = .4; ReversalRatio = .6; MinCycles = 4). The second had fewer criteria with intermediate thresholds but a higher minimum number of cycles (PeriodConsistency = .6; AmplitudeConsistency = .6; MonotonicityInAmplitde = .6; FlankConsistency = .6; MinCycles = 5). The third set had fewer criteria but stricter monotonicity thresholds (frequency in range of narrowband filter; PeriodConsistency = .7; FlankConsistency = .3; MontonocityInAmplitude = .9; MinCycles = 3). These criteria were chosen a-priori based on manual tuning of the burst detection on an independent dataset of wake EEG in adults during sleep deprivation.

After bursts were detected in each channel separately, they were grouped into clusters when they occurred simultaneously in multiple channels with roughly the same frequency. The frequency of bursts was calculated as the inverse of the average distance between negative peaks (1/period). Bursts for which the shorter one overlapped at least 50% of the longer one, and were within 1 Hz of each other, were considered part of the same burst cluster. Bursts identified separately in each channel were used for all the topographies, otherwise burst clusters were used to reduce the effect of burst globality (spread across the scalp) on measures of density.

#### 2.3.1 Oscillation measures

*Oscillation amplitudes* were calculated as the average negative-to-positive peak voltage difference for all cycles involved in all bursts, with units in microvolts (μV). *Oscillation densities* were calculated as the percentage of the recording occupied by bursts (sum of all the bursts’ durations divided by the duration of the recording). This accounts for both the duration and overall quantity of bursts in the recording. The average duration and number of bursts can be affected by the background aperiodic signal which can break up sustained oscillations into smaller bursts (Tal et al., 2020), therefore burst density was preferable. When calculating across multiple channels (e.g. Figure 2), oscillation density could easily exceed 100%, as burst clusters in different frequency ranges often co-occur. When combining densities across multiple frequency bands, bursts were pooled rather than averaged (sum of the durations of all the bursts of any frequency, divided by the duration of the recording). In our previous publication (Snipes et al., 2023), we referred to oscillation densities as “quantities”, however this term did not properly account for the normalization in time.

#### 2.3.2 The choice of frequency bands

Only bursts between 2 and 16 Hz were detected. Below 4 Hz very few bursts could be identified, therefore only bursts above 4 Hz were included in the analysis. Bursts over 16 Hz could be detected, but with higher false-positive rates, as determined by visual inspection. The choice of cutoff at 16 Hz was done arbitrarily a-priori to capture alpha (8-12 Hz) with generous padding. The division of bands for Figure 6 and Figure 7 was done using conventional bands (Pernet et al., 2020) with 1 Hz gaps to reduce overlapping information due to the drift in peak frequencies across individuals and ages: theta (4-7 Hz), alpha (8-11 Hz) and low beta (12-16 Hz). The inclusion of low beta was done aposteriori based on results observed in Figure 5B.

Many researchers advocate for the use of an individual alpha frequency (IAF) to define frequency ranges, especially when analyzing data across development (Bazanova & Vernon, 2014; Klimesch, 1999; Ostlund et al., 2022; Tröndle et al., 2022). This is done by selecting the peak alpha frequency separately for each individual and defining the band around this peak. The shift in IAF with age makes a strong case for such an approach (Freschl et al., 2022). The main problem with using IAF is that it assumes the largest oscillation will be functionally the same for all participants. Given that our participants displayed large heterogeneity in the number and amplitude of frequency peaks within the 2-16 Hz range (Suppl. Figure 1-1), and the vast majority of our participants were old enough that the peak alpha frequency was larger than 8 Hz (Freschl et al., 2022), we preferred to use fixed bands with gaps. The completely distinct topographies across the three bands for all age groups support this decision for this dataset (Figure 6, Figure 7).

### 2.4 Spectral power analysis

Spectral power was calculated using MATLAB’s *pwelch* function, with 4 s Hanning windows and 50% overlap. When average power across channels was calculated, edge channels were excluded (total count: 98). To dissociate periodic and aperiodic spectral power, we used the MATLAB extension of specparam (formerly known as FOOOF (Donoghue et al., 2020)). Spectra were smoothed over 2 Hz, and the aperiodic signal was fitted between 2 and 35 Hz (frequencies sufficiently separated from the 0.5-40 Hz filter range of the preprocessed data). Otherwise, the default settings were used (peak width: [0.5 12], max number of peaks: inf; minimum peak height: 0; peak threshold: 2; aperiodic mode: fixed).

#### 2.4.1 Spectral and aperiodic measures

*Power* was calculated by averaging the log-transformed power values between 4 and 16 Hz. Aperiodic *offsets* were provided by specparam as the log power value at 1 Hz of the aperiodic signal, and *exponents* as the x value in the 1/*f*^*x*^ model that describes the steepness of the aperiodic signal. The values are such that positive exponents refer to a downward descending aperiodic signal, and the larger the value the steeper the descent. *Periodic power* was calculated as the log-transformed power from which the aperiodic signal was subtracted. Fitting model parameters *r-squared* and *mean absolute error* (MAE) were similarly analyzed to evaluate whether the model fit could account for the results (Ostlund et al., 2022), with the results provided in Suppl. Figure 2-1. Overall, the specparam model fit was very good, with average MAE of 0.028 (interquartile range: 0.019, 0.035) and r-squared values of 0.998 (0.997, 0.999).

### 2.5 Sleep EEG

A subset of wake data was compared to previously analyzed sleep data (Jaramillo et al., 2020), comprising the *sleep-wake dataset*. This included only neurotypical participants from Dataset2016 and Dataset2017, resulting in 44 participants, 50% female, with mean age of 16.3 (8.1-24.8). Only one night per participant (experimental baseline) was included. The corresponding wake data was the auditory oddball for Dataset2016 (>18-year-olds), and the TAP Alertness task for Dataset2017 (<18-year-olds). The channel-averaged data is provided in Suppl. Data 2.

EEG sleep data was recorded at 500 Hz with Cz reference. Sleep stages were scored according to standard AASM scoring guidelines (Iber, 2007). Epochs containing artefacts were rejected after visual inspection using a semi-automatic approach (Huber et al., 2000). Channels with poor signal quality were removed and interpolated, and outer edge channels were removed (total count: 98).

Traditionally, sleep pressure is quantified with average spectral power (“slow wave activity” is power between 0.5-4 Hz). However, directly measuring individual slow wave parameters is considered a more precise marker of sleep pressure and synaptic plasticity (Esser et al., 2007; Riedner et al., 2007; Vyazovskiy et al., 2009). This is why both here and in our previous study (Jaramillo et al., 2020) we analyze sleep data focusing on slow wave amplitudes and especially slopes.

Slow wave detection was performed similarly to Riedner et al. (2007). The EEG was bandpass filtered from 0.5 to 4 Hz (stopband 0.1 and 10 Hz, Chebyshev Type II filter) and rereferenced to the average. Negative deflections between zero-crossings were identified as slow waves if they were separated by 0.25-1 s. *Slow wave amplitude* consisted of the most negative amplitude between zero-crossings of the signal. The descending *slow wave slope* was the amplitude of the negative peak divided by the time from the positive-to-negative zero-crossing to the negative peak. The negative peak was used because this corresponded to silent periods of neuronal spiking, and the descending slope is thought to reflect the synchronization of this silent period across neurons (Nir et al., 2011; Vyazovskiy et al., 2009).

Slow waves were detected in the first and last hour of artifact-free NREM sleep. Given that slopes are directly calculated from amplitudes, slow waves from the first to last hour were first matched by amplitude (see Jaramillo et al. (2020) for details), and only the slopes of matched slow waves were included in the analysis (∼60% of total waves). Thus, the decrease in slow wave slopes across the night is independent from the decrease in amplitudes.

### 2.6 Statistics

Statistics were performed using the MATLAB Statistics and Machine Learning Toolbox. For all analyses, statistical significance was determined for p-values < .05. Given the heterogeneous datasets pooled together for this analysis, we chose to conduct linear mixed effects models to model the relationship between age, sleep, ADHD, and EEG measures. This was done with the function *fitlme*.

For each fixed factor of the model, β estimates, t-values, p-values, and degrees of freedom (df) are reported in the text. β estimates of continuous variables (e.g., age) indicate by how much the EEG measure (e.g. density) changes for each unit of the continuous variable (e.g., 1 year) when all other variables are 0. Similarly, the β estimates of categorical variables (e.g. group) indicate how much the EEG measure changes from that category (e.g., ADHD) to the baseline category (e.g., controls), for all other factors set to 0. Lastly, t-values allow a comparison of the magnitude of the effect of each factor when comparing models with different measuring units. To determine the topography of the effects, we plotted β estimates and their associated statistical significance, corrected for multiple comparisons across channels using false discovery rates (FDR; (Benjamini & Yekutieli, 2001)).

We ran models with fixed factors Task, Time, Age, Group, and Sex, the interaction between Time and Age, and nested mixed factors Participant and Session. *Task* compared the levels *Oddball* vs. *go/no-go*, *alertness*, and *fixation*. *Time* compared the time of recording; evening vs. morning for wake, and first vs. last hour of NREM sleep. *Group* compared neurotypical participants vs. those with ADHD. *Sex* compared females vs. males. Depending on the analysis and subset of the data, different factors were included or not, and so the exact model is specified before each analysis and in each figure caption.

As a simpler analysis and sanity check, we conducted Pearson’s correlations between age and each wake measure, including only auditory oddball recordings, averaging sessions (Figure 2). In each figure, r values are provided as effect sizes. Pearson’s correlations were also conducted between all measures in the sleep-wake dataset (Suppl. Figure 2-2).

## 3 RESULTS

### 3.1 Effects of age, sleep, sex, and ADHD on wake EEG measures

The following linear mixed effects model was applied to each wake EEG measure, averaged across channels: Measure ∼ Task + Time ∗ Age + Group + Sex + (1|Participant) + (1|Participant: Session). The full outputs of the model are provided in Suppl. Data 3.

Oscillation amplitudes significantly decreased with age (β = -0.783, t = -11.33, p < .001, df = 1234), significantly decreased the morning following sleep (β = -3.977, t = -15.90, p < .001, df = 1234), with a significant positive interaction (β = 0.143, t = 7.74, p < .001, df = 1234), such that amplitudes decreased less overnight with increasing age (Figure 2, bottom row). Amplitudes were significantly lower in males than females (β = -1.476, t = - 2.38, p = .017, df = 1234), and were not significantly different in participants with ADHD (β = -0.574, t = -0.89, p = .373, df = 1234). As can be seen in Figure 2, the relationship between age and amplitudes was quite robust, both as absolute values (r_eve_ = -.66, r_mor_ = -.63) and overnight changes (r = .56). Overall, amplitudes changed in the expected directions for both development and sleep pressure, except for the lack of an effect of ADHD.

Oscillation densities significantly decreased with age (β = -3.540, t = -2.11, p = .035, df = 1234), and the morning after sleep (β = -77.506, t = -10.39, p < .001, df = 1234), with a significant positive interaction between age and time of recording (β = 4.878, t = 8.85, p < .001, df = 1234). Unlike amplitudes, the correlation between age and density was weak (r_eve_ = -.28, r_mor_ = .01). Instead, the correlation between age and overnight change in density was quite strong (r = .57), such that oscillation densities decreased overnight in children under 15 and increased in young adults (Figure 2). There was no effect of ADHD (eta = -0.154, t = -0.01, p = .992, df = 1234) or sex (β = -16.446, t = -1.10, p = .271, df = 1234). Overall, oscillation densities behaved independently from amplitudes, especially in the direction of overnight changes in adolescents and adults.

Aperiodic exponents became significantly shallower with age (β = -0.027, t = -9.23, p < .001, df = 1234) but significantly steeper overnight (β = 0.099, t = 5.51, p < .001, df = 1234), with a trending negative interaction between time of recording and age (β = - 0.003, t = -1.91, p = .056, df = 1234). The increased overnight steepness was driven primarily by a decrease in higher frequency power (Suppl. Figure 5-1), whereas decreasing steepness with age is driven by decreases in low frequency power (Favaro et al., 2023). The correlations between exponents and age were as robust as for oscillation amplitudes (r_eve_ = -.60, r_mor_ = -.66), but the correlation with overnight change was weak (r = - .15, not significant). There was no significant effect of ADHD (β = 0.028, t = 1.07, p = .284, df = 1234) or sex (β = 0.041, t = 1.63, p = .102, df = 1234). This means that aperiodic exponents also change independently from oscillation amplitudes, and do not reflect the direction of changes expected for sleep pressure.

Aperiodic offsets also significantly decreased with age (β = -0.067, t = -16.59, p < .001, df = 1234), but with no significant effect of time (β = 0.029, t = 1.65, p = .100, df = 1234), and a significant negative interaction (β = -0.004, t = -3.11, p = .002, df = 1234), such that offsets decreased more overnight with age. The correlations between age and offsets were the strongest of all measures (r_eve_ = -.81, r_mor_ = -.83), however, the correlation between age and overnight change in offset was negligible (r = -.05). Again, there was no effect of ADHD (β = 0.021, t = 0.55, p = .579, df = 1234) or sex (β = -0.003, t = -0.07, p = .944, df = 1234). Overall, offsets correlated with age in the same direction as amplitudes, changed overnight in the same direction, but the overnight change was not larger in children compared with adults, counter to what would be expected for sleep pressure.

Curiously, fitting parameters of the specparam algorithm showed both significant age and time of recording effects, with the model fit improving in the morning and worsening with age (Suppl. Figure 2-1). Therefore, differences in fit quality may partially contribute to the effects observed for aperiodic measures. However, the changes in model fitting go in the opposite direction one would expect of data quality (i.e. worse in the morning and better in adults). This suggests that some other aspect of the EEG signal that changes with age and time can affect the fit quality of the specparam algorithm.

In summary, of the four EEG measures, only amplitudes followed the same trajectories expected for both development and sleep pressure. The absolute values of all four measures had a negative correlation with age, and differed primarily in the overnight response and the relationship between age and overnight response. Oscillation densities in particular showed a strong effect of age on overnight changes, reversing direction between childhood and adolescence. No measure showed any relationship with ADHD, and only amplitudes were affected by sex. Therefore, in later analyses we did not include these factors, and pooled patients and controls for greater statistical power.

#### 3.1.1 Comparison of oscillation and aperiodic measures to spectral measures

To quantify the extent to which wake spectral measures were influenced by oscillation and aperiodic measures, we directly correlated each measure to the other in the sleepwake dataset (Suppl. Figure 2-2), and then controlled for time of the recording, task, and age using linear mixed effects models in the complete wake dataset: Measure1 ∼ Measure2 + Time ∗ Age + Task + (1|Participant) + (1|Participant: Session) (Suppl. Table 2-1).

Without correcting for anything, all measures significantly correlated with each other, except periodic power which did not correlate with either aperiodic exponents or offsets. Power was most related to amplitudes, also in the mixed effects model (correlation r = .93, mixed effects t-value = 39.3). Power was also significantly related to densities (r = .82, t = 28.2) and offsets (r = .80, t = 17.5), but less so with exponents (r = .54, t = 4.7). Periodic power was most correlated with densities (r = .80, t = 46.9) and somewhat with amplitudes (r = .53, t = 20.6).

#### 3.1.2 Comparisons between wake and sleep EEG measures

Using the sleep-wake dataset, we directly compared wake measures with sleep slow wave amplitudes and slopes, first through simple correlations (Suppl. Figure 2-2) and then with the linear mixed effects model: SleepMeasure ∼ WakeMeasure + Time ∗ Age + Task + (1|Participant). The full models are provided in Suppl. Data 4.

All wake measures were significantly correlated with both slow wave amplitudes (wake amplitude r = .61; density r = .27; exponent r = .46; offset r = .66) and slopes (amplitude r = .70; density r = .38; exponent r = .60; offset r = .80). When controlling for the time of the recording, task, and age, wake amplitudes were significantly related to both slow wave slopes (β = 4.21, t = 2.8, p = .006, df = 82) as well as slow wave amplitudes (β = 0.78, t = 2.7, p = .007, df = 82). The same was true for wake power (slow wave slopes: β = 26.36, t = 2.3, p = .023, df = 82; slow wave amplitudes: 5.84, t = 2.6, p = .011, df = 82). While aperiodic offsets had shown the strongest correlations to sleep measures, with the mixed effects model they were only significantly related to amplitudes (β = 14.69, t = 2.8, p = .006, df = 82) but not slopes (β = 40.26, t = 1.6, p = .111, df = 82). No other wake measure was related to either slopes (density: β = 0.05, t = 1.1, p = .289, df = 82; exponent: β = 7.78, t = 0.3, p = .785, df = 82; periodic power: β = 44.92, t = 1.2, p = .247, df = 82) or amplitudes (density: β = 0.02, t = 1.2, p = .235, df = 82; exponent: β = 5.99, t = 0.8, p = .453, df = 82; periodic power: β = 7.69, t = 0.8, p = .412, df = 82).

Overall, wake amplitudes were most related to sleep slow waves, with significant relationships to both amplitudes and slopes. This association was reflected in wake power. Wake offsets were also significantly correlated to sleep measures, however the association between offsets and sleep slopes did not survive the correction for age and time in the mixed effects model.

### 3.2 Topography of wake EEG measures by age

Figure 3 provides the average topographical maps of each measure for five age bins, averaging (or pooling for densities) all frequencies from 4-16 Hz, from the oddball task. Amplitudes, densities, exponents, and offsets all showed unique topographies from each other. Across ages, for each measure there were primarily changes in magnitude more so than major regional differences. However, oscillation amplitudes in the youngest cohort began as a single midline occipital spot, which spread bilaterally in the 7–10-year-olds. Prominent central bilateral peaks also appeared in the 7–10-year-olds. Oscillation densities similarly started as a single midline occipital spot, but these became more lateral-parietal in the 14–18-year-olds. Like amplitudes, two small bilateral central peaks emerged in the densities of 7–10-year-olds, which merged with the primary occipital-parietal cluster in the 14–18-year-olds. Furthermore, a frontal peak gradually emerged with age. Exponents were steepest in midline channels, whereas offsets showed both a frontal midline and occipital peak. As with the correlations between measures, power topographies most resemble amplitudes, and periodic power resembles densities.

**Figure 3:**
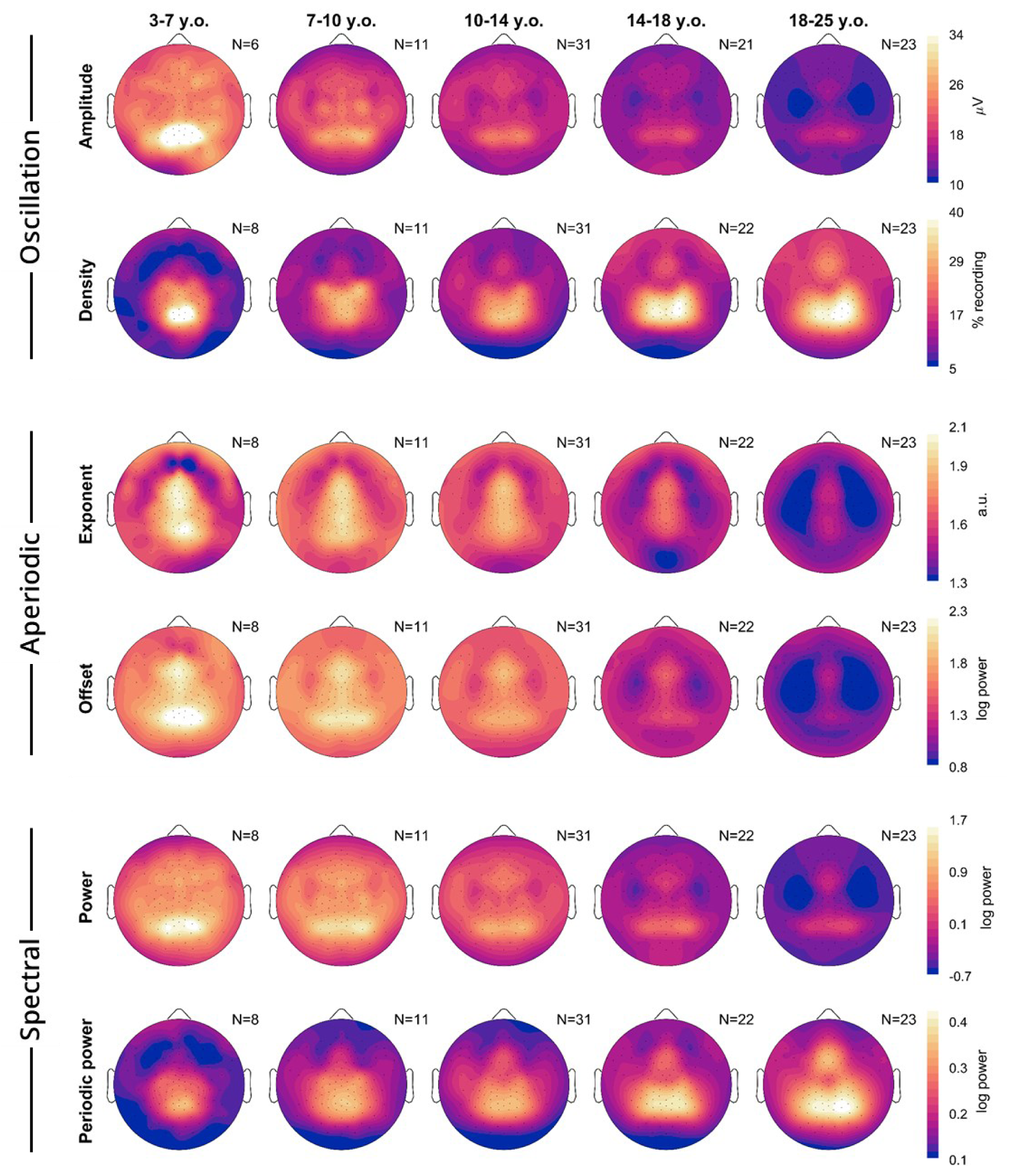
Topography averages of wake EEG measures. Each plot is a schematic of the EEG viewed from above, with the nose on top. Lighter colors indicate greater magnitude over a given location for that measure (rows). Only neurotypical participants and oddball recordings were included, and participants were grouped into age bins (columns). Multiple oddball recordings from different sessions and times of day were first averaged for each participant. The number of participants included is indicated in the top right corner of each plot. Acronyms: y.o., years-old; a.u., arbitrary units.

To determine the topography of overnight changes in EEG measures, we performed linear mixed effects models for each channel, with the model Measure ∼ Time + Task + (1|Participant) + (1|Participant: Session), dividing participants into 4 age bins (the youngest 3-7 were excluded as they were too few, with too few recordings). Figure 4 plots the β estimates for the effect of Time.

**Figure 4:**
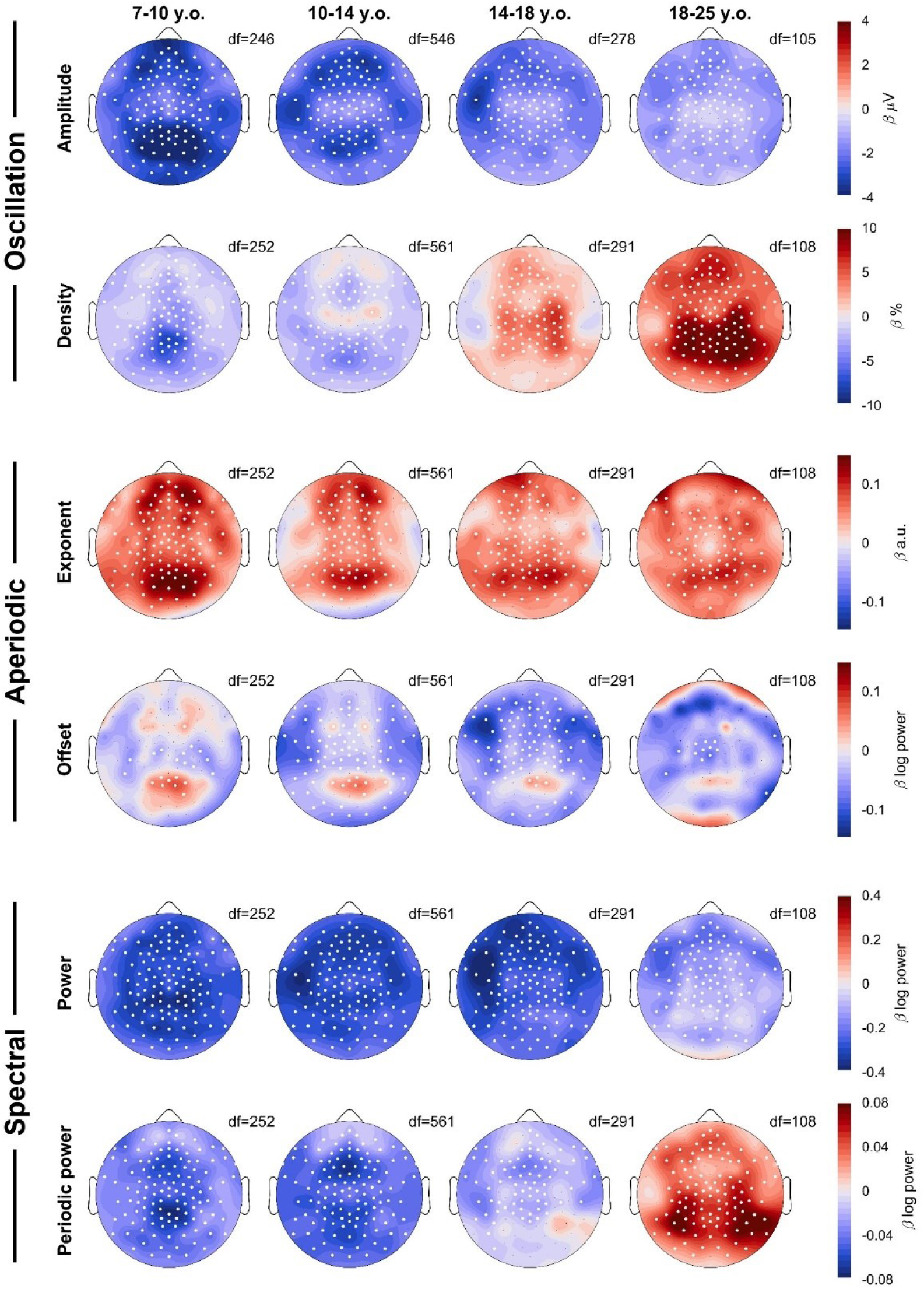
Topographies of wake EEG overnight changes. A linear mixed effects model was run for each measure, each age group, and each channel: Measure ∼ Time + Task + (1|Participant) + (1|Participant:Session). Color reflects β estimates for the fixed effect *Time*, such that red indicates an overnight increase in that measure. The factor *Task* was not included for the 18-25 y.o. group, as these participants only performed oddballs. White dots indicate channels for which the β estimate was statistically significant, corrected for multiple comparisons with FDR (false discovery rate). Black dots indicate remaining channels. Data includes both patients and neurotypical controls. Degrees of freedom (df) are provided for each plot. For each column, the sample sizes were N = 29, 67, 36, & 23.

Amplitudes showed widespread overnight decreases across all age groups, however the decrease was largest in occipital channels for the youngest group, and slightly more fronto-temporal in young adults. The overnight density topographies resembled the average density topographies from Figure 3, in terms of location of the effects. The youngest group showed the largest overnight decrease in the same midline-occipital spot where there were the most oscillations (Figure 3), and adults showed the largest increase in the same bilateral occipital-parietal areas where they had the largest densities.

The overnight increase in the steepness of exponents was widespread, but peaked in an occipital spot in all age groups, with additional bilateral frontal spots in <18-year-olds. These topographies do not correspond to the average topography of exponents from Figure 3. Offsets revealed widespread decreases, with localized increases in the same occipital locations for which exponents increased the most. This suggests that aperiodic offsets generally decrease, although the increase in exponent steepness contrasts this effect.

Power and periodic power again showed similarities to amplitudes and densities respectively, however while densities increased in the 14–18-year-olds, periodic power decreased. Likewise, the overnight increase in periodic power for >18-year-olds was more occipital and lateral than the increase in densities, and the larger decrease in amplitudes in occipital regions of 7–10-year-olds was less evident in power than for amplitudes.

### 3.3 Frequency-specific wake EEG changes with age and sleep pressure

In the previous analyses, we had pooled all frequencies between 4 and 16 Hz. Subsequently, we explored how oscillation and spectral EEG measures changed for each frequency, this time averaging channels. Average evening values for each age and frequency are plotted in Figure 5A, and the overnight differences are plotted in Figure 5B.

**Figure 5:**
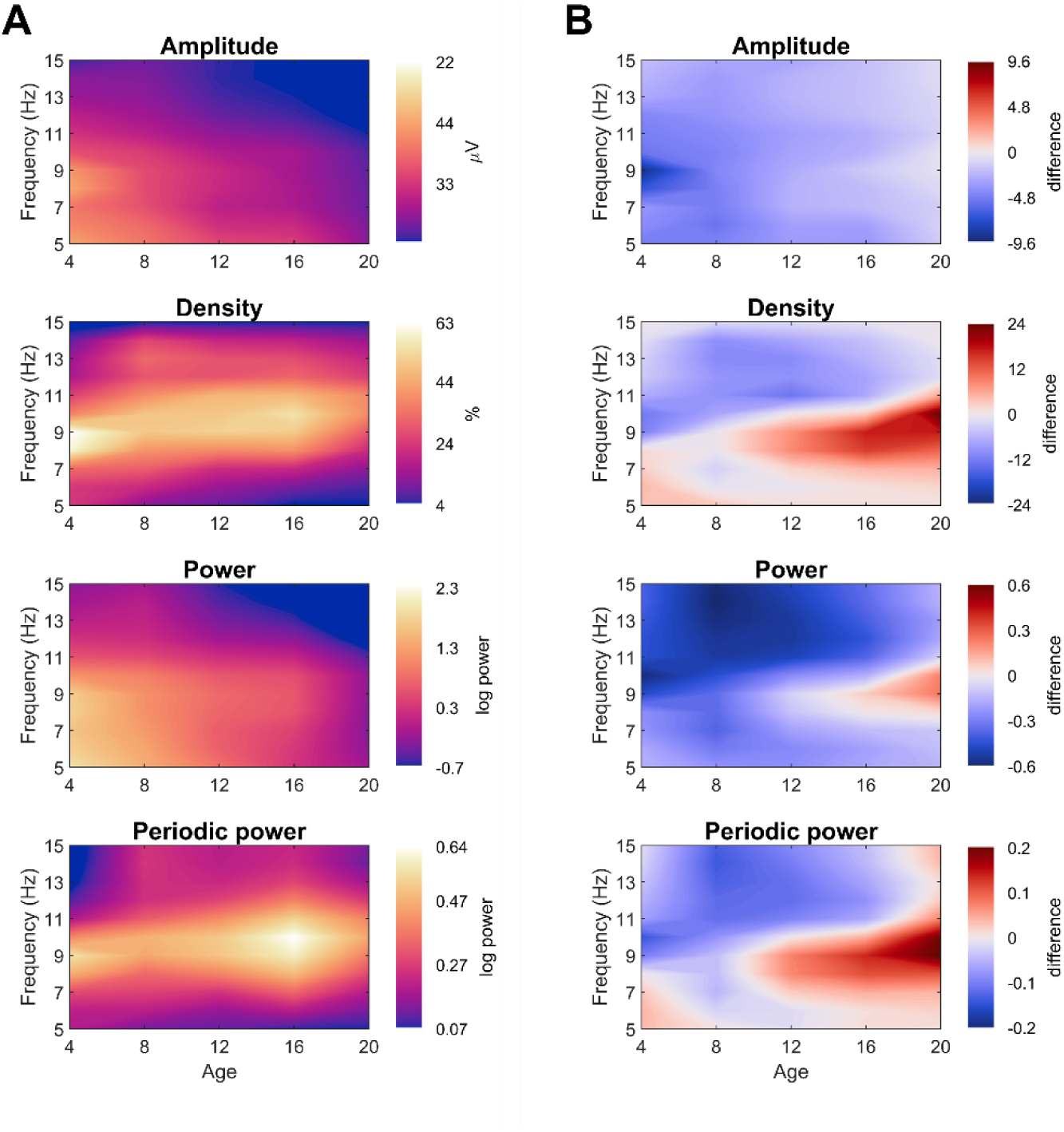
Average spectrograms across ages. From the oddball task, pooling controls and ADHD participants. Average spectra from the other tasks are provided in Suppl. Figure 5-1. **A**: Average values, such that lighter colors indicate greater magnitude for a given frequency and age. For spectra extending to older ages and sleep stages, see Sun et al. (2023). **B**: Difference values between morning and evening recordings, such that red indicate a greater magnitude in the morning. The measurement unit of each figure is the same as that of A.

Average oscillation amplitudes and spectral power showed gradual gradients, with highest values in the youngest participants and lowest frequencies, and lowest values in the oldest participants and highest frequencies. Like average amplitudes, overnight changes in amplitude showed largely gradual decreases with age and frequency.

Average density and periodic power instead had distinct peak values between 8 and 11 Hz, with the peak shifting upwards with age, a well-known property of alpha oscillations during development (Freschl et al., 2022; Smith, 1938; Tröndle et al., 2022). Overnight densities (and periodic power) showed decreases in higher frequencies (>11 Hz, i.e. low beta) and increases in alpha. These increases only began between 8-10 years of age, they were strongest in adults, and the range shifted to higher frequencies with age.

Given the dissociation between alpha and beta for density, we explored the topographic changes in density, split by both age and frequency. In Figure 6, we plot average values. Theta oscillations were the overall rarest. They were most prevalent in the youngest children, and with age, the peak in theta density gradually shifted upward and decreased in magnitude. Alpha density instead started as a midline occipital cluster in the 3–7-year-olds. Alpha densities in the occipital spot decreased in the 7-10 cohort, with bilateral central spots instead becoming more pronounced. With age, these three peaks morphed into a continuous occipital-parietal cluster, while in the remaining channels overall density of alpha increased. Lastly, low beta oscillations showed yet another topography. They were almost completely absent in the youngest group, and started to appear in the 7–10-year-olds as lateral occipital peaks and a frontal midline spot. Gradually, the lateral peaks converge towards the midline, and in adults became partially overlapping with alpha, on average right-lateralized.

**Figure 6:**
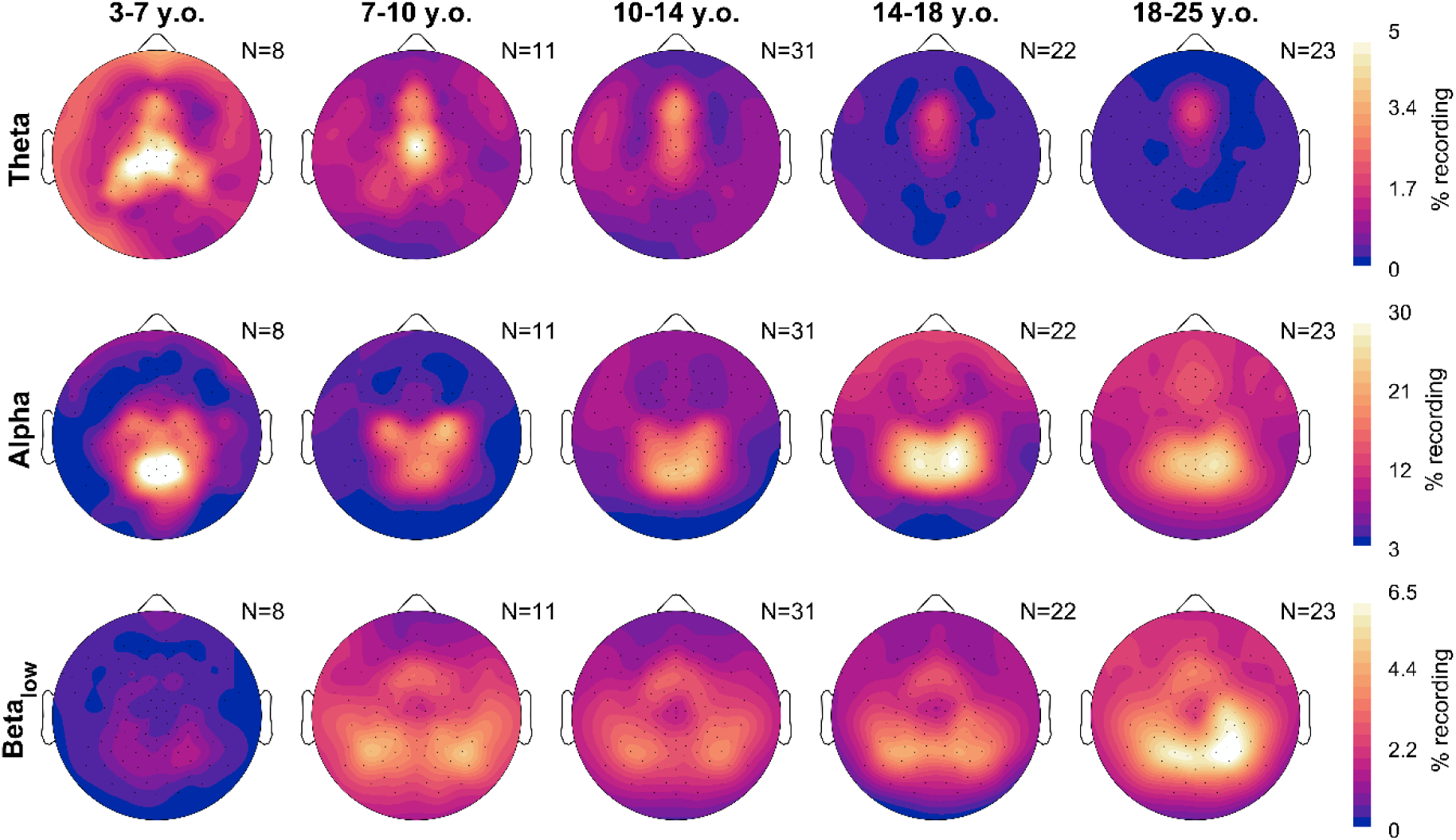
Average topographies of oscillation densities, split by frequency band. Recordings are the same as in Figure 3. Theta is 4-7 Hz, alpha is 8-11 Hz, and low beta is 12-16 Hz. Each band has a different color scale range.

The overnight changes in density split by frequency are in Figure 7, based on the model Density ∼ Time + Task + (1|Participant) + (1|Participant: Session). Like in Figure 4, these reflect the β estimates for the fixed effect of Time.

The overnight changes in theta density were small (around 1%), however quite variable by region and age. In the 7–10-year-olds, theta density generally increased overnight except in a central spot, exactly where the largest theta densities were seen in Figure 6, which instead decreased. This continued for the 10–14-year-olds. In the 14–18-year-olds, there were no significant effects, however the midline theta spot, now more frontal, showed a decrease. In adults, only some scattered theta increases were observed.

**Figure 7:**
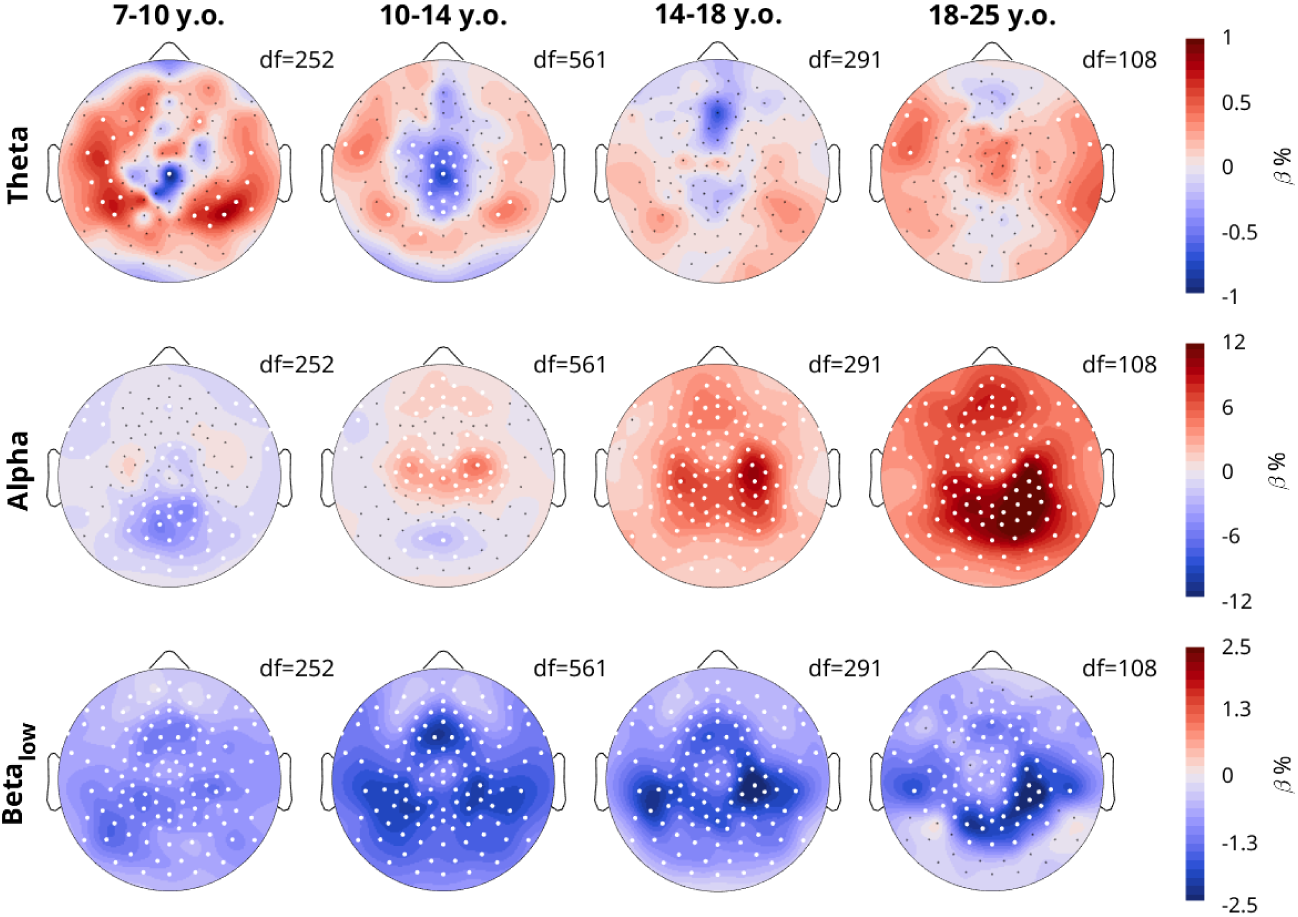
Topographies of overnight density changes, split by frequency band. Participants and plot information are the same as in Figure 4, with color indicating the β estimates for the linear mixed effects model, such that red indicates an overnight increase in density. The model was Density ∼ Time + Task + (1|Participant) + (1|Participant:Session).

For alpha, the main occipital spot in the 7–10-year-olds decreased overnight. Already in the 10–14-year-olds a bilateral central alpha rhythm started to increase overnight, with still some slight decreases in the occipital spot. The overnight increases spread to the entire scalp in adolescents and adults, peaking in occipital parietal areas, especially right lateralized. For low beta, across all ages there were decreases, with the peak shifting across age bins.

### 3.4 Effect of ADHD on the wake EEG topography

In our initial mixed effects models pooling channels and frequencies, we found no significant effects for ADHD, which is why in subsequent analyses and figures we no longer included a Group effect. However, we nevertheless conducted mixed effects models to determine the effect of ADHD for each channel: Measure ∼ Task + Time ∗ Age + Group + (1|Participant) + (1|Participant: Session). We found no significant effects when correcting for multiple comparisons (Figure 8). However, amplitudes were on average lower in participants with ADHD and frontal exponents were steeper compared to controls.

**Figure 8:**
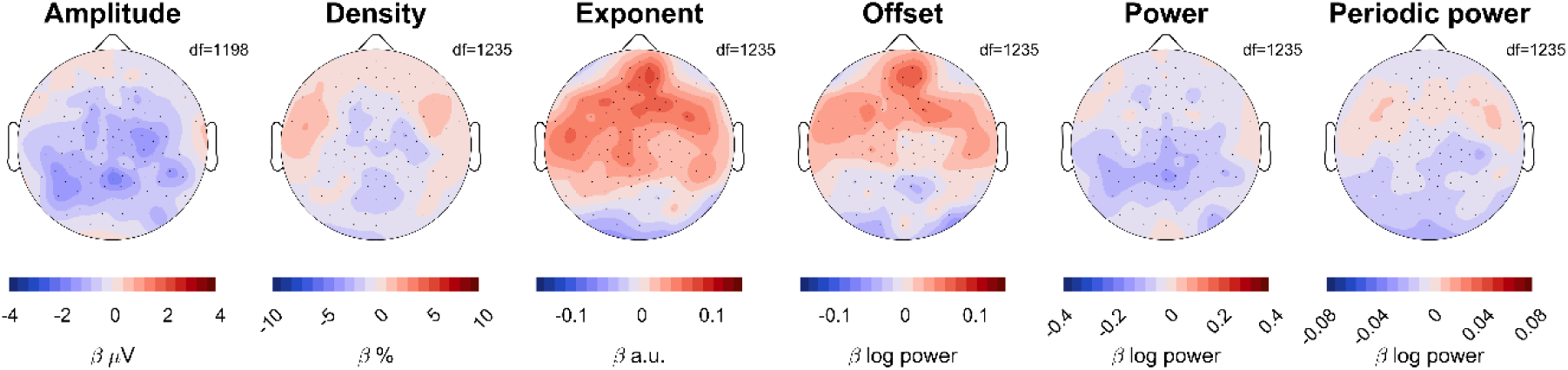
Effects of ADHD on EEG measures. Red indicates larger values in patients compared to controls. The scale for each topography is the same as for Figure 4. The model was Measure ∼ Time*Age + Task + Group + (1|Participant) + (1|Participant:Session). White dots would have indicated statistically significant channels, following FDR correction for multiple comparisons.

## 4 DISCUSSION

In this study, we compared four distinct wake EEG measures and their relationship to development, sleep pressure, and ADHD. Specifically, we hypothesized that oscillation amplitudes would behave like sleep slow waves, because both measures likely reflect neuronal synchronization due to synaptic density and plasticity. Our predictions were met on all accounts except for the sensitivity of wake amplitudes to ADHD. Of the four measures, only amplitudes decreased overnight in all ages *and* the overnight decrease was largest in younger children (Figure 2). Wake amplitudes were significantly correlated to sleep slow wave measures, also when correcting for age and participant. Aperiodic offsets showed similar development and overnight changes, as well as strong correlations to sleep measures, but they did not display larger overnight decreases in younger children, a key indicator of higher neural plasticity.

While oscillation amplitudes were the only wake measure that followed all the patterns expected for sleep pressure, every EEG measure reflected development. Average amplitudes, exponents, and offsets were all strongly anticorrelated with age, with offsets showing the strongest relationship (Figure 2). The correlation between average densities and age was weak, but we found distinctive regional patterns across age groups (Figure 6). Overnight changes in both oscillation amplitudes and density were robustly correlated with age (Figure 2, Figure 7), whereas overnight changes in exponents and offsets were not as affected by age, as evidenced by the low correlations in Figure 2. These results are summarized in Figure 9. Finally, no measure showed significant effects of ADHD (Figure 8).

**Figure 9:**
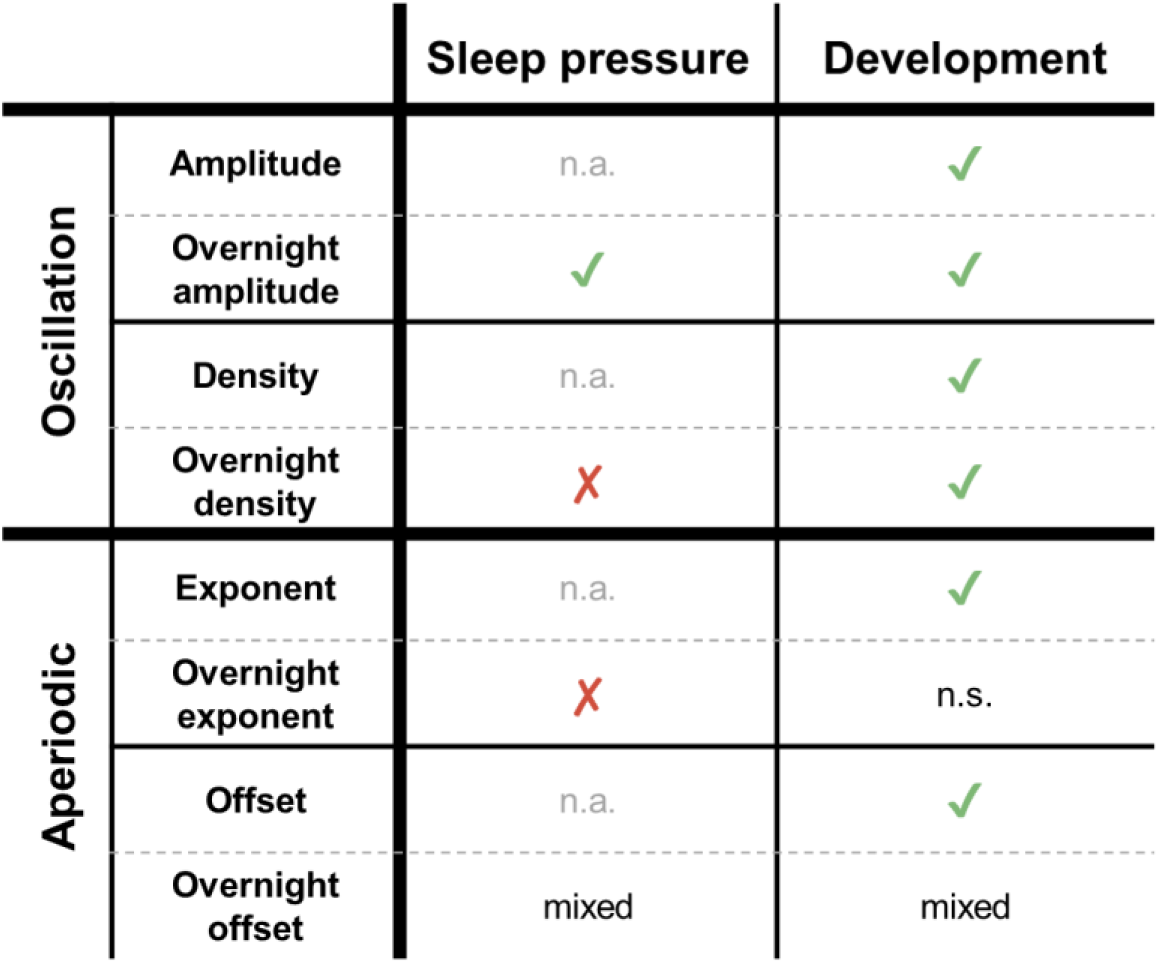
Summary of which measure reflects sleep pressure or development. Sleep pressure was determined by whether a measure decreased overnight and whether the decrease was larger in younger children. Development was determined by whether there were strong effects of age. Overnight offsets had “mixed” results across analyses. Acronyms: n.a., not applicable; n.s., not significant.

### 4.1 Oscillation amplitudes

Like for slow waves during sleep, the decrease in wake oscillation amplitudes with age could be explained by decreasing synaptic density in the cortex across adolescence (Huttenlocher, 1979). The overnight decrease in wake amplitudes could be explained by sleep’s role in reducing net synaptic strength (Cirelli & Tononi, 2022; Tononi & Cirelli, 2014), and such plastic changes are more pronounced in children than adults (Jaramillo et al., 2020). The larger decrease in occipital regions in children compared to adults may be because primary sensory and motor areas obtain peak cortical thickness earlier in children, followed by adjacent secondary areas and finally frontal association areas (Shaw et al., 2008), resulting in an overall posterior-anterior maturation trajectory (Kurth et al., 2010). Therefore, younger children show larger overnight decreases in amplitudes in occipital areas because these areas undergo higher plastic changes at that maturational stage. Finally, even the significant sex difference was comparable to sleep slow wave activity (Mourtazaev et al., 1995), with wake oscillation amplitudes higher in females than males (although the effect was relatively small).

Higher slow wave activity in females has been hypothesized to reflect their smaller heads and thinner skulls (Dijk et al., 1989). It is then also possible that head and skull size affect the age-related changes we observed for wake amplitudes. However, Schaworonkow and Voytek (2021) did not find any changes in wake oscillation amplitudes from 1-7 month-old infants, despite this period corresponding to the fastest growth in head circumference (Roche et al., 1986). Therefore, it is unlikely that the more modest increases in head size from childhood to adolescence can entirely explain the large decrease in amplitudes we observed here.

While wake oscillation amplitudes behave largely like sleep slow waves, the effects are reduced. In adults the overnight decrease in wake amplitudes was near 0 μV (Figure 2), which is not the case for either slow wave slopes or amplitudes during sleep (Jaramillo et al., 2020; Riedner et al., 2007). One possible explanation for this is that the evening wake recordings fell within the wake maintenance zone. This is a circadian time window just before bedtime characterized by increased alertness (Shekleton et al., 2013; Strogatz et al., 1987). We had found in our previous study that oscillation amplitudes are significantly reduced in this window, counteracting the otherwise monotonic buildup in amplitudes that occurred throughout the day (Snipes et al., 2023). In adults the contrasting effect of the wake maintenance zone may be sufficient to equalize evening and morning amplitudes. Overall, wake oscillation amplitudes may reflect the same information as sleep slow waves but may be more affected by circadian or other factors. Therefore, wake amplitudes may be affected by the same underlying neuronal changes as sleep slow wave activity, but they are not sufficiently specific nor sensitive to replace this gold standard measure of sleep pressure.

### 4.2 Oscillation densities

As could be expected, the densities of oscillations were the most complex EEG measure, with effects differing depending on age, time, topography, and oscillation frequency.

Theta oscillations were most prevalent in early childhood, and decreased progressively with age, supporting previous results measuring relative theta power (Somsen et al., 1997). We further found that the peak in theta densities steadily drifted more frontally across childhood and adolescence. This frontal theta in adults is known to originate from the midline prefrontal cortex, and to be anti-correlated to the default mode network (Ishii et al., 2014; Michels et al., 2010; Scheeringa et al., 2008). Therefore, this drift in theta may reflect the steady maturation of both frontal cortices and the default mode network (Fan et al., 2021).

There is still unresolved contradictory evidence on the role of theta in adults (Snipes et al., 2022), without including the question of theta during development. On the one hand, theta is often associated with cognitive effort (Buzsáki, 2005; Cavanagh & Frank, 2014; Meyer et al., 2019; Mitchell et al., 2008). On the other, it is also associated with sleepiness (Finelli et al., 2000; Smith, 1938; Snipes et al., 2022) and fatigue (Arnau et al., 2021; Tran et al., 2020; Wascher et al., 2014). A possible resolution to this paradox is that there are distinct oscillations that originate from different circuits with different functions and just happen to occur at the same frequency. Our results in Figure 7 would support this, as the peak source of theta shows overnight *decreases*, whereas theta from the rest of the cortex instead shows overnight increases, even as the theta peak drifts more frontally. Alternatively, theta could reflect a general form of “idling rhythm” (Snipes et al., 2022, 2024), originating from disengaged cortical areas, and what changes from evening to morning is which circuits tend to idle. Theta as an idling rhythm is supported by simultaneous EEG-fMRI studies that find theta activity anti-correlating with brain metabolism (Scheeringa et al., 2008). This would make theta functionally comparable to alpha (Laufs et al., 2003), differing only by source and frequency.

Alpha begins in young childhood as a midline spot (Figure 6). Two lateral central peaks become more defined at 7-10 years of age. These likely reflect sensorimotor mu rhythms which appear when motor activity is absent or even suppressed (Pfurtscheller et al., 2006; Pineda, 2005), and is already present in infants (Berchicci et al., 2011). We find that with age, they become topographically indistinguishable from occipital alpha, at least when recorded during an oddball task. These lateral central peaks are the first to show overnight increases in 10-14 year-olds, while the overnight decrease in the occipital midline spot becomes less prominent. The overnight increase in alpha densities then spreads over bilateral parietal and occipital areas across adolescence and adulthood. This dissociation between overnight decreases in childhood and increases in adulthood, as well as the slight differences in topography, could suggest that occipital alpha is in fact qualitatively distinct in children and adults. However, these rhythms were previously considered functionally equivalent because also in infants alpha power increases with eyes closed compared to eyes open (Stroganova et al., 1999). More research is needed on the sources of these oscillations.

It is also possible that this dissociation in overnight density changes is driven by some other difference with age, such as a longer window of sleep inertia in young children, longer sleep duration, or a shifted circadian rhythm compared to adults. Melatonin in the morning is elevated in children under 10, whereas older children and adolescents have morning melatonin levels comparable with the rest of the day (Attanasio et al., 1985). In adults, alpha power fluctuates with circadian rhythm and is therefore synchronized to melatonin levels (Cajochen et al., 2002). Therefore, it is possible that the dissociation of decreasing/increasing alpha densities originates from children and adults being at different phases of their alpha circadian rhythm in the morning. More research is needed into the circadian effects on the EEG across development.

Finally, we found overnight decreases in low beta densities across ages. There is ample literature on “beta bursts” from sensorimotor areas (Feingold et al., 2015; Jones et al., 2009; Little et al., 2019; Lundqvist et al., 2024; Shin et al., 2017; Wessel, 2020), however these are likely qualitatively distinct from the beta oscillations we observed. First, the topography of sensorimotor beta rhythms was found to be either bilateral-central in infants or frontal in adults (Rayson et al., 2023), unlike the bilateral occipital topographies in Figure 6. Second, sensorimotor beta was found to occur in “bursts” of power less than 150 ms long (Sherman et al., 2016) and could thus correspond to only 1-3 cycles of beta (Jones, 2016; van Ede et al., 2018). Supporting this, lagged coherence decreases rapidly after 2 cycles (Rayson et al., 2023). We detected bursts that were *at least* 3 cycles long, only up to 16 Hz, and therefore always longer than 187 ms. Therefore, the low-beta bursts we captured with cycle-by-cycle analysis are likely distinct from the previously investigated sensorimotor beta activity. It remains an open question what is the functional role of these low-beta oscillations, and why they may be less common the morning after sleep.

### 4.3 Aperiodic offsets and exponents

Our results on offsets and exponents replicate previous findings: they decrease linearly with age (Cellier et al., 2021; Hill et al., 2022; Tröndle et al., 2022) and originate from broad, primarily midline sources (Favaro et al., 2023). As in Lendner et al. (2023), we found exponents becoming increasingly steeper after sleep, extending this finding across ages. We further found, unusually, that exponents and offsets change in opposite directions, with offsets decreasing after a night of sleep. For aperiodic activity measured *during* sleep, both exponents and offsets decrease across the night, and the decrease decreases with age (Horváth et al., 2022). This indicates that changes in aperiodic activity related to sleep-wake history do not reflect the same information when measured during wake or during sleep. It is possible that during sleep the decrease in exponents reflects the decrease in sleep pressure, whereas during wake the increase in exponents could reflect something like lingering sleep inertia in the morning. Notably, the change in aperiodic exponents with age and sleep depth affect primarily lower frequencies (Favaro et al., 2023), whereas the overnight wake EEG changes affect primarily higher frequencies (Suppl. Figure 5-1). This means that the pivot point (the frequency at which the aperiodic signal rotates) of the exponent differs, and this likely has biological significance. Future studies should, as in Bódizs et al. (2021), specifically identify such pivot points, also to better dissociate changes in offset and exponent.

Second to oscillation amplitudes, aperiodic offsets showed the closest resemblance to sleep slow wave activity, decreasing overnight and showing strong correlations with slow wave measures. While aperiodic offsets were not significantly related to slow wave slopes when controlling for age and participant, this does not imply a significant difference from wake oscillation amplitudes. However, offsets did not show the characteristic decrease in overnight changes with age that was observed for slow waves and wake amplitudes (Figure 2). This could suggest that offsets also reflect changes in synaptic strength and plasticity, but may be less sensitive or specific than wake amplitudes, especially to overnight changes.

### 4.4 Spectral power vs oscillation burst detection

As explained by previous papers, changes in spectral power do not differentiate between changes in oscillation amplitudes or densities (Donoghue et al., 2022; Quinn et al., 2019; Snipes et al., 2023; Tal et al., 2020; Zich et al., 2020). With this study, we again demonstrate that amplitudes and densities meaningfully change independently, both across sleep and development. However, we also found that the changes in amplitude were largely captured by changes in average power, whereas changes in density were largely captured by changes in periodic power. This suggests that power and periodic power can be used as proxies for amplitudes and densities. Previous studies of development have likewise found differences between power and *relative power* (similar to periodic power) across development (Somsen et al., 1997). These results may be reinterpreted as the dissociation between changes in amplitude and density.

Arguing against this approach, however, is the topography of overnight changes in 14– 18-year-olds: oscillation densities increased overnight, whereas periodic power decreased, likely reflecting the greater influence of the overnight decrease in amplitudes. Similarly, the overnight increase in densities is more central than the overnight increase in periodic power. Therefore, differences between power and periodic power may suggest different influences of oscillation amplitudes and densities, but to know for certain, oscillations should be measured directly.

### 4.5 ADHD

Despite a relatively largesample size (N = 58), we did not observe any significant effects of ADHD on our EEG measures. We did find exponents to be steeper on average in patients, supporting the results of Robertson et al. (2019), and contrasting those of Ostlund et al. (2021). One explanation could be that our participants were performing tasks for most recordings, and the differences between patients and controls may mostly emerge in resting EEG; patients may have developed compensation mechanisms masking potential differences during the tasks. It’s also possible our analysis did not reach significance because our participants were a combination of both medicated and unmedicated patients, and it is known that medication will reduce the effects of ADHD on the EEG (Furrer et al., 2019; Karalunas et al., 2022). Additionally, our participants were screened for good sleep quality (to have a chance of falling asleep in the laboratory with an EEG net, and to have similar levels of sleep pressure as controls). However, around 40-55% of children with ADHD report sleep deficits (Becker et al., 2019; Corkum et al., 1998; Holmberg & Hjern, 2006; Konofal et al., 2010), so it is possible that poor sleep quality in patients contributes to differences in the wake EEG observed in prior studies (Clarke et al., 2020). Finally, it’s possible that only subtypes of patients with ADHD have a specific relationship to any of the measured EEG markers; ADHD is highly heterogeneous with varying symptoms among individuals.

Regardless of the reason, given that we do not see any systematic differences between patients and controls, none of the wake EEG measures we tested make for a reliable intrinsic marker of ADHD which could potentially be used to aid diagnosis. Instead, research investigating such markers in these and other patient populations should take special care to control for sleep/wake history and sleep quality, as these may have a greater impact on the EEG. Instead, within the same participants we observed significant differences in sleep slow wave activity from controls (Furrer et al., 2019), suggesting sleep is more sensitive to developmental deficits.

### 4.6 Limitations

An important limitation of this study is the scarcity of datasets under 8. It is known that in young children there is a switch from primarily synaptic growth to primarily synaptic pruning, peaking in different cell populations and regions at different ages (Cao et al., 2020; Petanjek et al., 2011; Shaw et al., 2008), which is also reflected in sleep slow wave activity peaking around this age (Feinberg & Campbell, 2013). Wilkinson et al. (2024) found an increase with age in both offsets and exponents across infants 0-3 years old, and McSweeney et al. (2023) found a quadratic relationship between age and offsets/exponents, such that they peaked at 5-7 years old. This would suggest more complex relationships between age and EEG measures than the linear trends observed here.

A further limitation of the data is its lack of uniformity, pooling multiple experimental paradigms and tasks, and medicated and unmedicated patients. Our primary findings remain significant also for more uniform subsets of the data (e.g. Figure 2 is only oddball, Suppl. Figure 2-2 only neurotypical controls), but it’s possible that different tasks would generate different topographies, and significant effects of ADHD would have emerged from entirely unmedicated patients.

This paper poses two potential weaknesses in interpretability. First, if one assumes that aperiodic and oscillatory signals are a linear sum of each other in the time-domain of the EEG signal, it is possible that the aperiodic changes explain part of the changes in oscillation amplitudes. A better understanding of the relationship between oscillations and the aperiodic signal are needed to validate such an assumption, and then precise simulations would be needed to determine the magnitude of this effect. However, in practice we have demonstrated that oscillation amplitudes can behave differently from aperiodic measures, and therefore it is at least useful to measure them independently. Second, overnight changes do not dissociate between homeostatic (related to sleep pressure) and circadian (related to clock time) effects. To do so would require substantially more intensive protocols, such as sleep deprivation, sleep restriction, or shifting sleep windows over several days. However, collecting more wake recordings throughout the day would already provide an indication as to when an effect is circadian or homeostatic.

Finally, our data is limited to EEG. Future studies and analyses would greatly benefit by comparing these measures to structural and functional brain changes observable with MRI, as well as cognitive and behavioral outcome measures related to development. This would bridge the gap between a purely basic research finding to practical applications.

### 4.7 Conclusions

We have found that overnight changes in wake oscillations provide distinct markers of brain maturation. Both absolute amplitudes and overnight changes in amplitudes decrease linearly with age, the effect more occipital in younger children. Overnight changes in oscillation density, especially of alpha oscillations, dissociate children from adolescents and adults by switching from an overnight decrease to an increase in density. Understanding the reason behind this effect would likely provide important information on brain development around puberty and adolescence. More generally, we have shown that there are a multitude of changes in the EEG with development that go beyond simple spectral power, each with their own functional significance. Moving forward, researchers should analyze independently oscillatory amplitudes and densities as well as aperiodic exponents and offsets, as they offer different insights into neuronal activity and structure.

## 5 DATA AND CODE AVAILABILITY

The data tables used for the linear mixed effects models are provided as supplementary material (Suppl. Data 1, Suppl. Data 2), and the output of the linear models for wake and sleep data are likewise provided (Suppl. Data 3, 4). The raw and preprocessed EEG is available upon request, requiring permission from the local ethics committee.

The burst detection can be conducted either with the original python toolbox bycycle (https://github.com/bycycle-tools/bycycle), or with our MATLAB implementation (https://github.com/HuberSleepLab/Matcycle). The preprocessing and analysis code is likewise open source (https://github.com/snipeso/children-wake/).

## 6 AUTHOR CONTRIBUTIONS

Methodology, SS, EK, VJ, CV, ML, SK, RH; Software, SS; Formal Analysis, SS, VJ; Investigation, EK, VJ, CV, MF, MS, ML, SK, RH; Resources, ML, OGJ, RH; Writing–Original draft, SS; Writing-Review & Editing, SS, EK, VJ, MF, MS, SK, OGJ, RH; Supervision, RH; Funding acquisition EK, ML, OGJ, RH

## 8 ACKNOWLEDGEMENTS & FUNDING

This work was supported by the Swiss National Science Foundation (PP00A-114923, PP00P3-135438, CRSII3_160803, 320030_153387, 320030_179443, PCEFP1_181279), the University Research Priority Program of the University of Zurich, Forschungszentrum für das Kind (FZK) of the University Children’s Hospital Zurich, the Clinical Research Priority Program “Sleep and Health” of the University of Zurich, the HMZ Flagship Project “SleepLoop” of University Medicine Zurich, the Borbély-Hess Foundation, Gottfried und Julia Bangerter-Rhyner-Stiftung, the Waterloo Foundation (2462-4978), and the National Institute of Health (K01MH074643). We thank everyone involved in data collection, and preprocessing, especially Maya Ringli, Renato Merki, Michelle Suppiger, Léa Geiger, Martina Liechti, Stefano Maurizio, Dr. Renate Drechsler, Stephanie Cares, Margot Schaerer, Jessica Bühler, Natalie Birnbaum, Demy Marcella Gutjahr, and Soraya Souissi. We thank the children and parents for their participation, and Giulio Tononi for the loan of technical material.

