## Supplementary material for "Wake EEG oscillation dynamics reflect both sleep pressure and brain maturation across childhood and adolescence": SupplMaterial

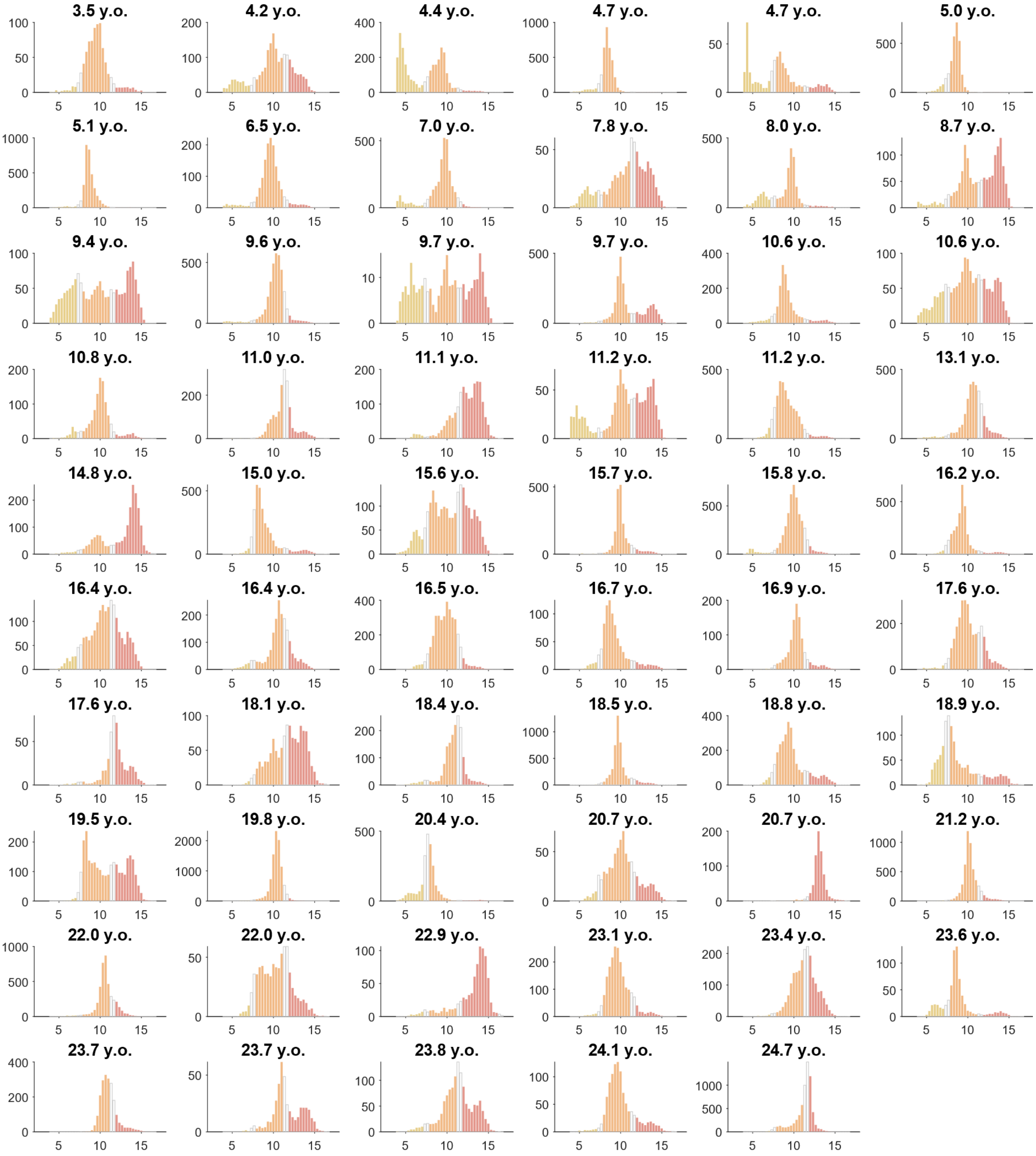


**Supplementary Figure 1-1: Individuals' distribution of oscillation densities by frequency.** Each plot is from one participant, with frequency (Hz) on the x-axis and density (% recording) on the y-axis. Colors reflect the frequency bands used in Main Figure 6 and Figure 7: yellow for theta (4-7 Hz); orange alpha (8-11 Hz); red low beta (12-16 Hz). This figure only includes evening oddball recordings from neurotypical participants, sorted by age.

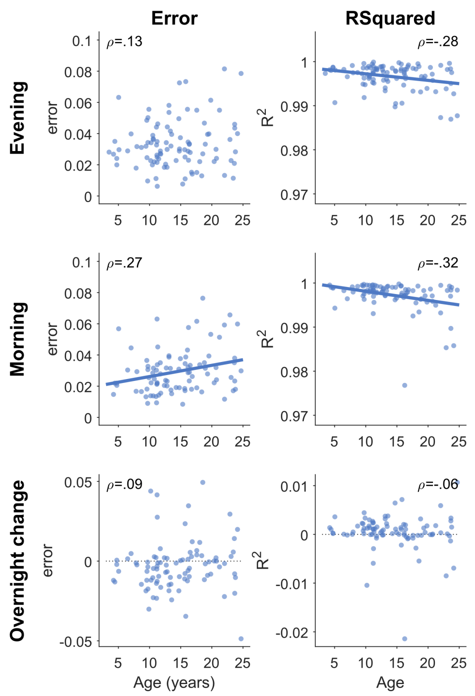


**Supplementary Figure 2-1: Specparam fitting estimates correlated with age.** Only auditory oddball recordings are included, pooling both neurotypical and ADHD participants, the same as in Main Figure 2. Each dot represents a single participant. For participants with multiple sessions, values across sessions were first averaged. Pearson's correlations were done for each figure, with rho values provided in the corner. If the p-value was less than .05, a correlation line was drawn (without correcting for multiple comparisons).

The same linear mixed effects model was applied to the fitting estimates of the specparam model, to evaluate whether any potential differences in model fitting could explain the aperiodic results. Mean absolute errors had a trending effect of age (beta = 0.000, t = 1.69, p = .091, df = 1234), curiously increasing with age, and a significant decrease the morning after sleep (beta = -0.007, t = -3.75, p < .001, df = 1234). There was no significant effect of ADHD (beta = 0.001, t = 0.36, p = .717, df = 1234) or sex (beta = 0.000, t = 0.09, p = .930, df = 1234), and the Time * Age interaction was only trending (beta = 0.000, t = 1.75, p = .080, df = 1234). R-squared values significantly decreased with age (beta = -0.000, t = -3.72, p < .001, df = 1234), significant increased after sleep (beta = 0.001, t = 2.96, p = .003, df = 1234), had no significant effect of ADHD (beta = -0.000, t = -0.87, p = .384, df = 1234), sex (beta = -0.000, t = -0.47, p = .639, df = 1234), or Time * Age interaction (beta = -0.000, t = -0.90, p = .366, df = 1234). Given that model fitting varied systematically with the factors of interest, it is possible some of the effects observed for aperiodic exponents and offsets are attributable to differences in model fitting. However, the main effects of aperiodic exponents and offsets were substantially larger than these effects of the model fits.

| | AMPLI-TUDE | DENSITY | EXPONENT | OFFSET | POWER | PERIODIC POWER |
|---|---|---|---|---|---|---|
| AMPLITUDE | | b=10.00<br>t=13.2<br>p<.001 | b=-0.00<br>t=-0.5<br>p=.618 | b=0.02<br>t=8.7<br>p<.001 | b=0.07<br>t=39.3<br>p<.001 | b=0.02<br>t=20.6<br>p<.001 |
| DENSITY | b=0.01<br>t=12.0<br>p<.001 | | b=0.00<br>t=14.7<br>p<.001 | b=0.00<br>t=14.5<br>p<.001 | b=0.00<br>t=28.2<br>p<.001 | b=0.00<br>t=46.9<br>p<.001 |
| EXPONENT | b=-0.76<br>t=-1.9<br>p=.063 | b=167.38<br>t=14.9<br>p<.001 | | b=0.88<br>t=61.9<br>p<.001 | b=0.18<br>t=4.7<br>p<.001 | b=0.17<br>t=12.5<br>p<.001 |
| OFFSET | b=3.05<br>t=7.6<br>p<.001 | b=160.06<br>t=14.5<br>p<.001 | b=0.84<br>t=61.0<br>p<.001 | | b=0.63<br>t=17.5<br>p<.001 | b=0.12<br>t=8.4<br>p<.001 |
| POWER | b=7.54<br>t=40.5<br>p<.001 | b=181.32<br>t=29.9<br>p<.001 | b=0.10<br>t=5.8<br>p<.001 | b=0.32<br>t=19.4<br>p<.001 | | b=0.22<br>t=30.2<br>p<.001 |
| PERIODIC POWER | b=13.77<br>t=19.3<br>p<.001 | b=666.33<br>t=46.7<br>p<.001 | b=0.61<br>t=12.1<br>p<.001 | b=0.46<br>t=8.5<br>p<.001 | b=1.73<br>t=27.9<br>p<.001 | |

**Supplementary Table 2-1: Mixed effects models between wake EEG measures.** The model was $Measure_{column} \sim Measure_{row} + \text{Task} + \text{Sleep*Age} + (1|\text{Participant}) + (1|\text{Participant:Session})$. The degrees of freedom for all models was df=1235.

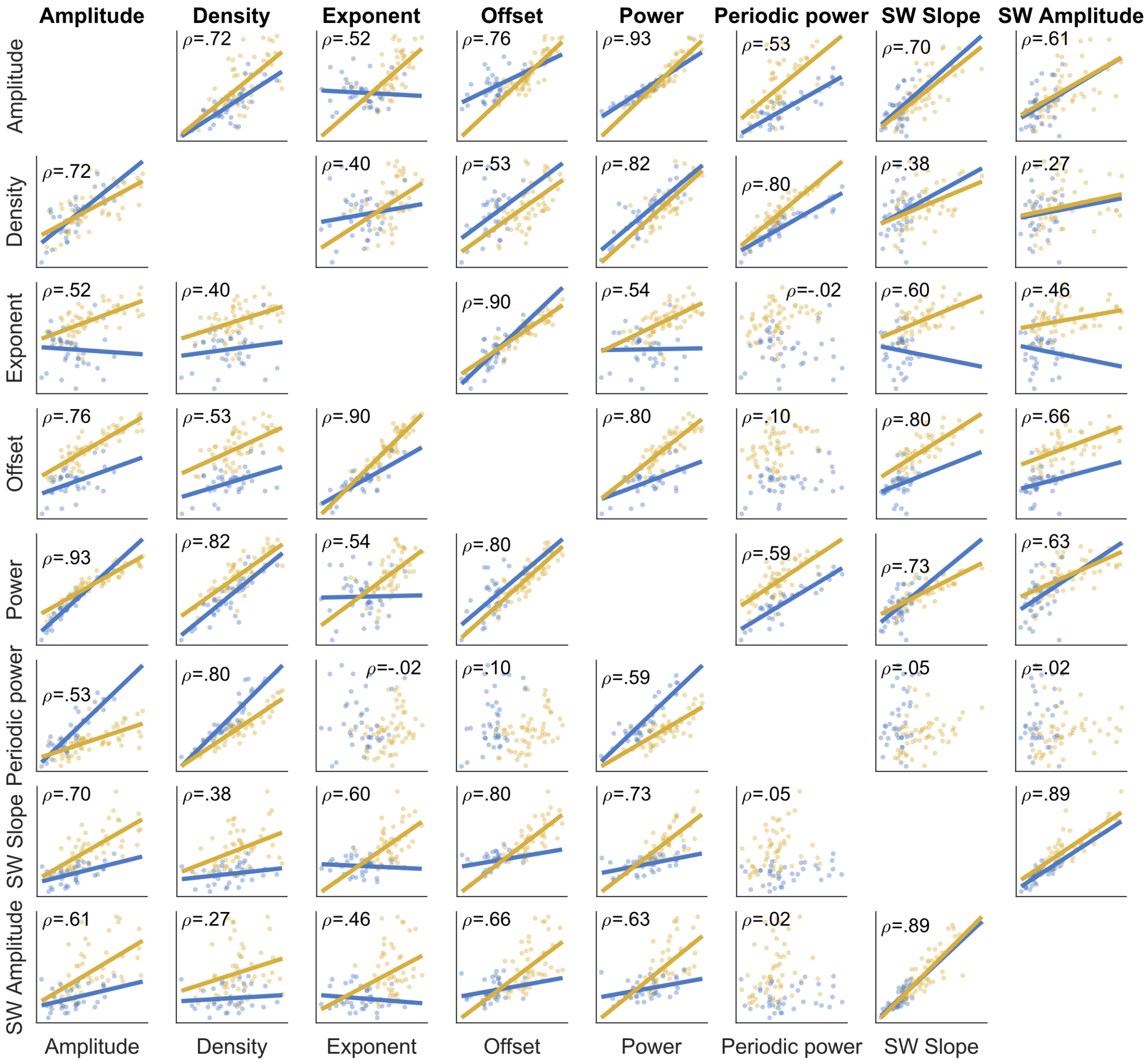


**Supplementary Figure 2-2: Correlations between outcome measures.** Yellow data is from Dataset2017 (<18 year olds, Alertness task) and blue data is from the Dataset2016 (>18 year olds, oddball task). Each dot represents the data of a recording for a single (baseline) session for each participant; each participant has both an evening and morning recording per plot. Sleep slow wave (SW) slopes and amplitudes were derived from the first or last hour of NREM sleep, correspondingly matched to either the evening or morning wake recordings. Pearson's ρ values are provided correlating all measures from both datasets, and significant correlations (p-value < .05) have linear fits plotted separately for each dataset. Given that there are repeated measures from the same participants (evening/morning) and two different wake datasets, these correlations violate assumptions of independence, but were conducted nevertheless to provide a simple metric to compare the relationship between different measures.

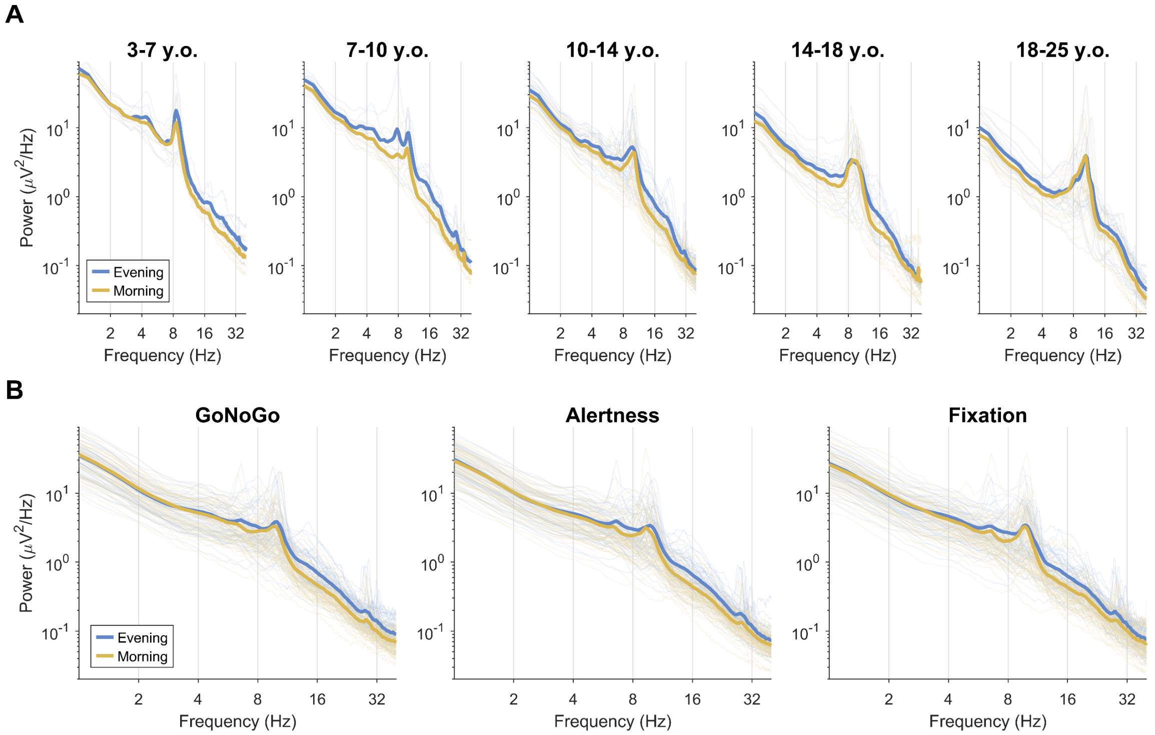


**Supplementary Figure 5-1: Spectrograms, averaged across channels.** Thick lines indicate group averages, thin lines indicate individuals (averaged across multiple recordings and channels), separately in the evening (blue) and morning (yellow). ADHD and controls are pooled. X- and y-axes are plotted on logarithmic scales. **A:** Auditory oddball tasks, split by age group, from datasets 2008, 2009, 2010, 2016. **B:** Go/no-go, alertness, and fixation recordings from datasets 2017 and 2019.
